# EnzOracle: Mechanism-aware prediction of enzyme environmental adaptation via a classification-guided mixture-of-experts framework

**DOI:** 10.64898/2026.06.02.729708

**Authors:** Qi Gao, Zhongcheng Fang, Yajing Yuan, Mengyuan Jin, Heqi Sun, Zhennan Peng, Lan Yang, Jiayi Li, Dong-Qing Wei

## Abstract

Industrial biocatalysis increasingly requires enzymes capable of operating under extreme physicochemical conditions, yet most natural sequence data reflect adaptation to mild environments, leading conventional predictive models to suffer from regression-to-the-mean effects in extremophilic regimes. Here we present EnzOracle, a classification-guided mixture-of-experts framework that enables distribution-aware prediction of enzyme melting temperature (*T_m_*), optimal catalytic temperature (*T_opt_*), and optimal pH (*pH_opt_*) directly from sequence. EnzOracle demonstrated robust predictive accuracy across diverse benchmarks, achieving RMSE of 5.245°C for *T_m_*, 11.458°C for *T_opt_*, and 0.781 for *pH_opt_*. Beyond predictive accuracy, we introduce a trait-resolved molecular simulation strategy to evaluate whether EnzOracle-derived attribution patterns correspond to independent physical mechanisms. Across representative systems, attention hotspots mapped onto rigidity-conferring interaction networks for *T_m_*, dynamically preorganized active-site ensembles for *T_opt_*, and pH-dependent electrostatic and hydration networks for *pH_opt_*. These orthogonal validations indicate that EnzOracle captures transferable biophysical principles of enzyme environmental adaptation rather than merely exploiting dataset-specific correlations, positioning sequence-based learning as a mechanism-aware framework for discovering stability and activity determinants across diverse catalytic landscapes.

## Introduction

Biocatalysis offers energy-efficient pathways for sustainable chemical synthesis; however, its industrial utility is strictly constrained by the physicochemical stability of native enzymes^1–6^. Consequently, the quantitative descriptors of enzyme fitness, specifically melting temperature (*T_m_*), optimal temperature (*T_opt_*), and optimal pH (*pH_opt_*), determine the viability of a biocatalyst for scale-up^7–9^. A critical disparity currently exists between genomic data availability and functional characterization. Although the UniProtKB database has expanded to approximately 200 million entries, experimental annotations for thermodynamic parameters remain sparse: fewer than 0.03% of these sequences have recorded *T_opt_* or *pH_opt_* values in the BRENDA database^10,11^. To bridge this sequence-function gap, deep learning (DL) has been adopted to predict physicochemical properties directly from amino acid sequences, acting as a computational filter to identify robust candidates from vast metagenomic libraries (Fig. S1a)^12–20^.

The paradigm of enzyme property prediction has shifted from empirical descriptors to data-driven representation learning. Early approaches relied on statistical aggregates of amino acid composition^21–24^. However, these shallow models struggled to resolve the nonlinear epistatic interactions governing protein fitness. The subsequent adoption of deep learning architectures enabled the automated extraction of local sequence motifs, a strategy exemplified by stability predictors^25–29^. More recently, self-supervised protein language models (pLMs) have further enhanced predictive accuracy by scaling learning to evolutionary magnitudes. Unlike shallow encodings, foundation models like ESM-2 project sequences into high-dimensional latent spaces that implicitly encode long-range structural constraints^30–32^. Benchmarking studies demonstrate that these evolutionary embeddings significantly outperform traditional physicochemical or one-hot encodings, providing the high-fidelity representations necessary for generalizing across diverse enzymatic families (Fig. S1b)^23,33,34^.

Despite these advances, a fundamental limitation remains: the misalignment between evolutionary data distributions and industrial requirements^35,36^. Natural evolution predominantly selects for fitness under physiological conditions, resulting in sequence databases that are heavily skewed toward mesophilic temperatures and neutral pH^37,38^. This distributional asymmetry poses a systemic hurdle for data-driven modeling. While heuristic strategies have been employed to mitigate this imbalance, they typically operate within the constraints of monolithic architectures^39^. By forcing a singular global model to simultaneously capture the distinct structural stabilization mechanisms governing both mesophilic and extremophilic behaviors, these methods often encounter conflicting gradients during optimization. Even with re-balancing, the gradients derived from the abundant canonical sequences tend to dominate the optimization trajectory, causing the model to treat high-value extremophilic features as statistical noise^24,40,41^. Addressing this architectural limitation requires moving beyond monolithic regression to context-aware frameworks capable of explicitly distinguishing between data-dense canonical regimes and data-scarce ecological niches prior to quantitative inference (Fig. S1c).

A second, equally critical challenge lies in the lack of mechanistic interpretability in sequence-based prediction models. While modern architectures achieve high predictive accuracy, it remains unclear whether model-derived features correspond to physically meaningful determinants of enzyme function, such as structural rigidity, catalytic preorganization, or electrostatic adaptation. This disconnect limits the utility of machine learning models for rational enzyme engineering, where mechanistic insight is essential for guiding mutation design.

To address these challenges, we present EnzOracle, a Classification-Guided Mixture-of-Experts (CG-MoE) framework that explicitly decouples the learning of data-dense canonical regimes from data-sparse extremophilic regimes. An adaptive gating network learns a continuous physicochemical routing manifold and dynamically integrates predictions from two specialized experts, thereby mitigating destructive gradient interference and enabling unbiased prediction across the full physicochemical spectrum. Importantly, we complement this architecture with a trait-resolved molecular simulation validation framework to test whether model attribution reflects physically interpretable mechanisms. By integrating attention-derived residue importance with atomistic molecular dynamics simulations, we demonstrate that EnzOracle-prioritized residues map onto distinct biophysical determinants, including rigidity networks governing thermal stability, near-attack conformations underlying catalytic efficiency, and electrostatic reorganization driving pH adaptation. This combined strategy establishes a direct link between sequence-level prediction and molecular mechanism, revealing that the model captures transferable biophysical principles rather than dataset-specific correlations.

Collectively, EnzOracle achieves highly reliable predictions for *T_m_*, *T_opt_*, and *pH_opt_* while simultaneously providing mechanistic interpretability, offering a unified framework for accurate prediction and rational design of industrial biocatalysts.

## Results

### The EnzOracle framework: a classification-guided paradigm for distribution-aware prediction

To address the optimization bias inherent in monolithic models, we developed EnzOracle, a deep learning framework based on a CG-MoE architecture (Fig. 1). Unlike conventional approaches that impose a uniform mapping across heterogeneous sequence spaces, EnzOracle explicitly decouples the learning of mapping rules for the majority distribution from the constraints of long-tail edge cases. The workflow operates through three hierarchically integrated stages: multi-modal representation learning, adaptive gating network, and conditional expert inference.

**Fig. 1.**
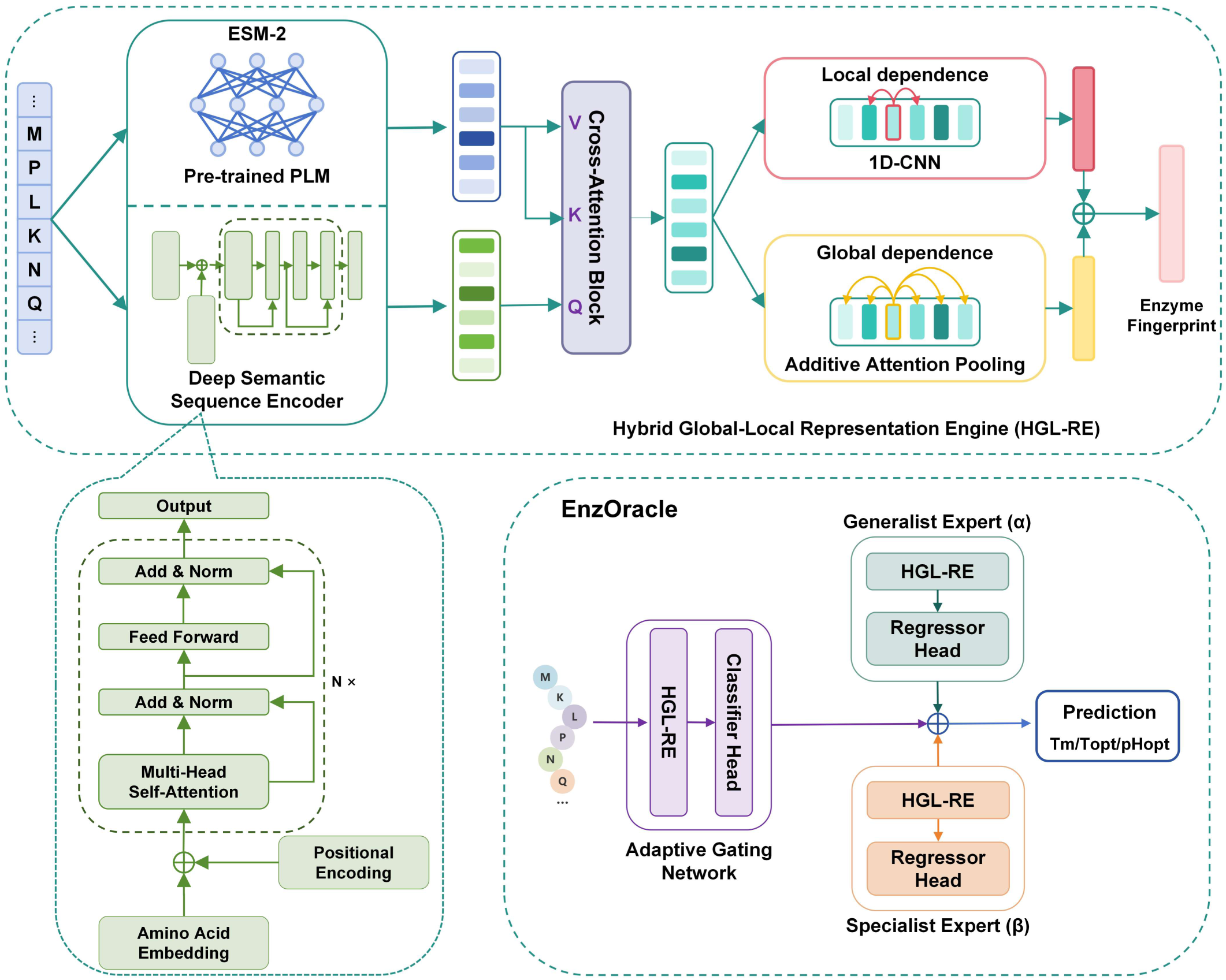
The overview of EnzOracle. The framework employs a Hybrid Global-Local Representation Engine (HGL-RE) to fuse evolutionary priors (from frozen ESM-2) with sequence features via Cross-Attention. An AGN functions as a gating network to route inputs within a fully decoupled Dual-Expert System. Unlike shared-parameter approaches, EnzOracle instantiates three independent HGL-RE encoders to prevent gradient interference, allowing the Generalist Expert (*α*) and Specialist Expert (*β*) to learn distinct feature distributions for the data-dense majority and data-sparse long-tail, respectively.

Effective functional prediction requires capturing both the local motifs of active sites and the global fold constraints of the protein. To achieve this, EnzOracle employs a Hybrid Global-Local Representation Engine (HGL-RE). Initially, raw amino acid sequences are encoded by a standard Transformer-based Sequence Encoder, which utilizes multi-head self-attention to capture long-range dependencies^42^. To compensate for data scarcity in functional datasets, we introduced an Evolutionary-Contextual Fusion Interface (ECFI). This module integrates representations from a frozen, pre-trained protein language model (ESM-2) as an external knowledge base. Through a cross-attention mechanism, the learned sequence embeddings serve as Queries (*Q*) to attend to the fixed evolutionary embeddings (Keys/Values, *K, V*), thereby enriching the representation with phylogenetic priors without requiring explicit multiple sequence alignments. The fused features are subsequently processed through two parallel pathways: a Global Attention Pathway (attention pooling) to extract holistic stability descriptors, and a Local Pathway (multi-layer 1D-convolutions) to capture spatially localized catalytic patterns.

A defining feature of EnzOracle is its use of conditional computation to mitigate gradient conflict. A dedicated Adaptive Gating Network (AGN) estimates the conditional probability of a query sequence belonging to the data-dense Generalist regime versus the data-sparse Specialist regime. This probability score serves as a dynamic weighting factor for a Dual-Expert System. Crucially, to prevent the dominant gradients of the majority distribution from diluting the feature representations of long-tail samples, we implemented a fully decoupled architecture. EnzOracle instantiates three independent HGL-RE encoders: one for the AGN, and two distinct encoders for the Generalist (*α*) and Specialist (*β*) experts. This design ensures that the high-dimensional embedding space of the Specialist expert is optimized exclusively for rare, edge-case constraints, with no parameter sharing or gradient leakage from the majority class. The final functional parameter is derived via a probability-weighted summation of the expert outputs, enabling smooth interpolation across the prediction manifold.

### EnzOracle enables robust and unbiased prediction of enzyme properties

To ensure a rigorous and direct comparison with established baselines, we utilized the official benchmark datasets from prior studies, preserving their original testing protocols. Statistical analysis confirmed that these evaluation sets for all three properties preserve the specific distributional profiles of the training corpus (Fig. S2). Critically, the testing sets effectively covered the extremophilic tails, retaining representative proportions of rare samples. Beyond distributional consistency, the benchmarks encompass a substantial proportion of sequences with low local homology to the training set (Fig. 2d-f), challenging the model to generalize beyond simple memorization. For *T_m_* prediction, the model achieved a coefficient of determination (*R*^2^) of 0.827 and a root-mean-square error (RMSE) of 5.245℃. Similarly, when compared to the experimental ground truth, predictions for *T_opt_* and *pH_opt_* yielded *R*^2^ values of 0.622 (RMSE=11.458℃) and 0.542 (RMSE=0.781 pH), respectively(Fig. 2a-c).

**Fig. 2.**
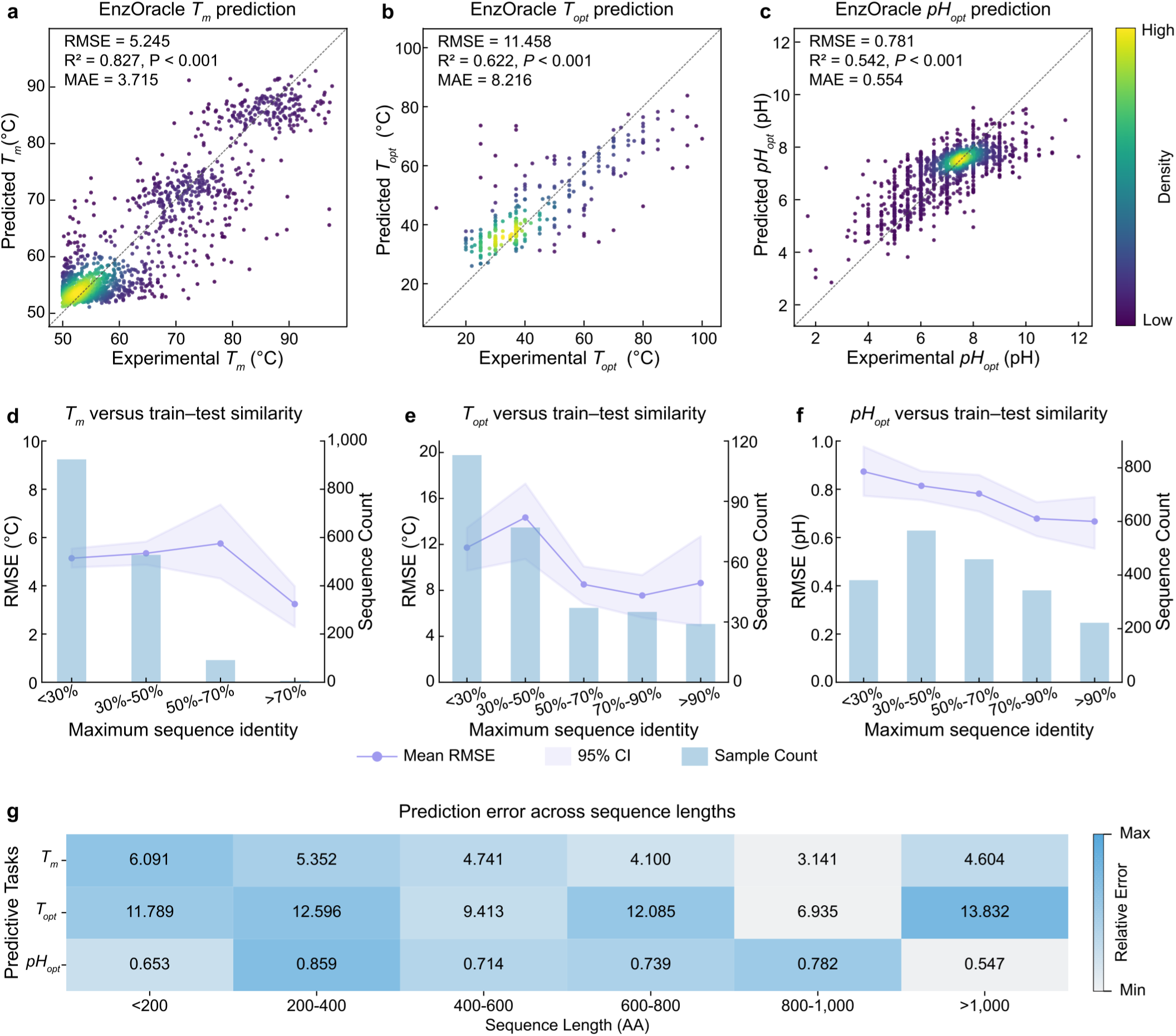
Global predictive performance and structural robustness of EnzOracle. **a-c**, Parity plots comparing predicted values against experimental ground truth for **a**, *T_m_*, **b**, *T_opt_*, and **c**, *pH_opt_*. Color intensity represents the local data density (kernel density estimation). The *R*^2^, RMSE, and mean absolute error (MAE) are denoted for each task. The dashed diagonal line indicates perfect agreement (*y*=*x*). **d-f**, Performance robustness across the sequence homology landscape. Test sequences were stratified by their maximum sequence identity to the training corpus (bins: <30%, 30-50%, 50-70%, 70-90%, >90%). The overlaid bar charts indicate the number of test sequences within each homology bin (right y-axis). Solid lines represent the mean RMSE, and shaded areas indicate the 95% confidence interval. **g**, Heatmap analysis of prediction error across varying structural scales. Rows represent sequence length intervals, and columns correspond to the three predictive tasks. Color intensity indicates the relative error magnitude (column-wise min-max normalized RMSE) to facilitate visual comparison within each task. The numbers annotated within the cells denote the raw RMSE.

A common failure mode in regression on imbalanced data is “regression to the mean,” where predictions collapse toward the dataset average. We investigated the source of prediction error by analyzing the residual distributions (Prediction-Truth). EnzOracle’s prediction errors exhibited symmetric, zero-centered profiles across all tasks (Fig. S3). The systematic bias (*μ*) was negligible, with *pH_opt_* achieving near-perfect alignment (*μ*=0.005 pH), significantly outperforming the thermal parameters (*T_m_*: −0.206℃; *T_opt_*: 0.784℃). Notably, the error distributions display pronounced leptokurtic characteristics (high peak at zero), distinctly deviating from standard Gaussian assumptions. This morphology indicates that the majority of predictions are highly concentrated around the ground truth with minimal deviation, while larger errors are confined to sparse outliers. The standard deviations (*σ*) of the residuals were controlled at 0.722 pH, 4.795℃, and 10.490℃, respectively. Collectively, the near-zero mean bias and the tight concentration of residuals confirm that EnzOracle is free from systematic drift and robust against the over-smoothing artifacts typical of monolithic models.

To assess whether EnzOracle learns universal physicochemical rules rather than merely memorizing homologous sequences, we evaluated its performance across the entire spectrum of sequence identities, confirming its robustness from highly homologous clusters down to the remote homology regime (<30% identity) (Fig. 2d-f). While *T_m_* and *pH_opt_* predictions remained remarkably invariant to sequence divergence, *T_opt_* prediction exhibited a characteristic dependency on evolutionary conservation, with RMSE increasing moderately in low-homology regimes. This distinction suggests that optimal temperature is more tightly coupled to specific, localized evolutionary motifs than global thermal stability. Additionally, we probed sensitivity to structural scale. Heatmap analysis (Fig. 2g) revealed that EnzOracle handles the full spectrum of protein sizes, from short peptides (<200 residues) to large complexes (>1000 residues). Notably, predictive accuracy for *T_m_* and *pH_opt_* actually improved for longer sequences (RMSE decreasing from 6.091°C to 4.604℃ for *T_m_*, and 0.653 to 0.547 for *pH_opt_*), suggesting that the Hybrid Global-Local Representation Engine effectively captures the dense long-range interaction networks that stabilize large folds.

Beyond sequence homology, we verified the model’s generalization across the biochemical functional landscape. We stratified the test set by Enzyme Commission (EC) numbers. EnzOracle exhibited negligible systematic bias across all seven catalytic classes, with prediction error distributions centered strictly around zero for both *T_opt_* and *pH_opt_* (Fig. S4).

### Bridging the accuracy-generalization gap across global and extremophilic physicochemical landscapes

To systematically evaluate predictive fidelity, we benchmarked EnzOracle against SOTA predictors. On global independent test sets, EnzOracle consistently established new performance benchmarks across all three tasks (Fig. 3a, c, e). For *T_m_* prediction, the model achieved an RMSE of 5.245°C, surpassing the leading baseline DeepSTABp (RMSE=5.368°C). The performance advantage was particularly pronounced in the complex *T_opt_* and *pH_opt_* tasks. EnzOracle reduced *T_opt_* prediction error to 11.458°C, significantly lower than the best competitor Seq2Topt (12.732°C), and achieved high-precision *pH_opt_* prediction (RMSE=0.781) compared to 0.845 for Catopt.

**Fig. 3.**
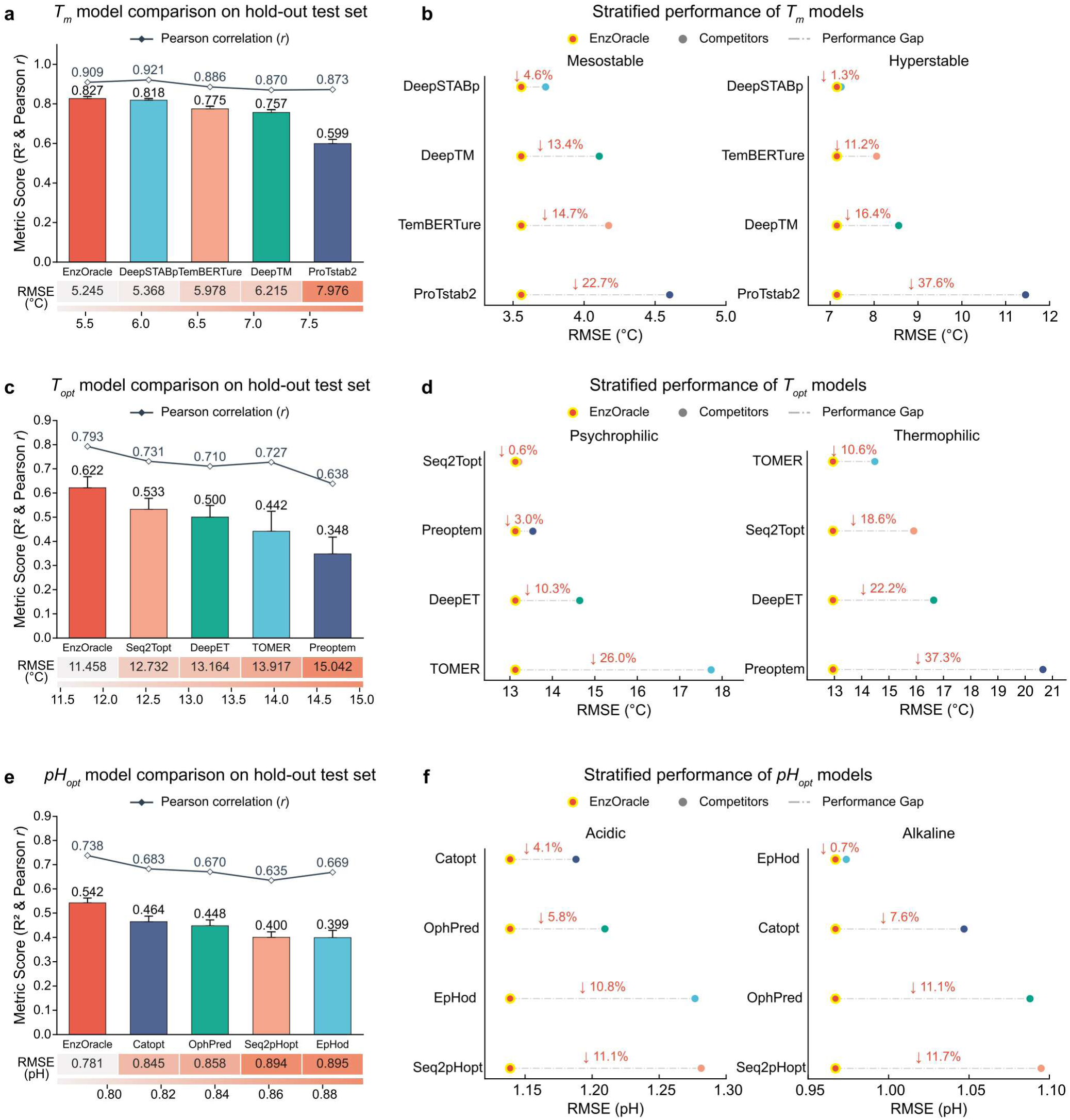
Bridging the accuracy-generalization gap across global and extremophilic regimes. **a**, **c**, **e**, Global benchmarking of predictive fidelity for **a**, *T_m_*, **c**, *T_opt_*, and **e**, *pH_opt_* on independent test sets. To provide a comprehensive assessment, bar heights represent the *R*^2^, the overlay line indicates Pearson correlation (*r*), and the color-coded heatmap below displays RMSE. **b**, **d**, **f**, Performance gap analysis in extremophilic environments. Pairwise comparison plots visualize the relative performance gain of EnzOracle against competitors in contrasting environmental regimes. The dashed lines and annotated percentages quantify the relative reduction in RMSE achieved by EnzOracle compared to specific competitors. **b**, Performance comparison for *T_m_* prediction in the mesostable (left, *T_m_*<60°*C*) and hyperstable (right, *T_m_*≥60°*C*) regimes. **d**, Performance comparison for *T_opt_* prediction in the psychrophilic (left, *T_opt_*≤30°*C*) and thermophilic (right, *T_opt_*≥50°*C*) regimes. **f**, Performance comparison for *pH_opt_* prediction in acidic (left, *pH_opt_*≤6.0) and alkaline (right, *pH_opt_*≥8.0) regimes.

While global metrics provide a broad overview, the critical bottleneck in biocatalysis lies in accurately characterizing enzymes in extremophilic regimes. We stratified the test datasets to quantify performance variations between canonical physiological environments and extreme physicochemical conditions. In all comparative analyses, prediction deltas were calculated against the top-performing baseline for each specific category to ensure a rigorous lower-bound estimate of performance gain. In the hyperstable regime (*T_m_*≥60°C), EnzOracle demonstrated superior robustness. Unlike baseline models such as DeepSTABp and DeepTM, which utilize auxiliary host Optimal Growth Temperature (OGT) as input to boost performance, EnzOracle (inferring solely from amino acid sequences) maintained a 1.3% lead over the top-performing DeepSTABp. Furthermore, it achieved a substantial 37.6% error reduction compared to ProTstab2 (Fig. 3b), validating the efficacy of the specialist-augmented architecture even without external data. Crucially, EnzOracle overcame the “regime specialization” limiting baseline models, particularly in *T_opt_* prediction. A prime example is the comparison with TOMER. While TOMER remains competitive in thermophilic conditions (*T_opt_*≥50°C) by leveraging host OGT information, its performance deteriorates significantly in the psychrophilic regime (*T_opt_*≤30°C). In contrast, EnzOracle demonstrates bidirectional stability, establishing a 26.0% lead over TOMER in the psychrophilic regime while simultaneously outperforming it by 10.6% in the thermophilic regime (Fig. 3d). Similarly, against the pure sequence-based method Seq2Topt, EnzOracle achieved relative error reductions of 0.6% (psychrophilic) and 18.6% (thermophilic). This broad-spectrum robustness was further confirmed in pH prediction, where EnzOracle outperformed the closest competitors by 4.1% in acidic (*pH*≤6) and 0.7% in alkaline (*pH*≥8) environments (Fig. 3f). These results indicate that EnzOracle effectively mitigates the accuracy-generalization trade-off, offering reliable predictions across the full physicochemical spectrum where conventional monolithic models often exhibit regime-dependent failure.

To assess the practical value of EnzOracle for virtual screening, we evaluated the “Screening Success Rate” (defined as the fraction of predictions falling within strict industrial tolerances) using REC analysis (Fig. S5)^43^. In identifying variants adapted to thermal extremes (*T_opt_*≤30°C or ≥50°C), EnzOracle attained a 33.2% success rate under a strict ±5°C tolerance Compared to the best sequence-based baseline success rate of 25.0%, this represents an absolute increase of 8.2%, corresponding to a substantial 32.8% relative performance gain. This screening advantage proved consistent in the predictive models for the other two physicochemical parameters.

Finally, we compared EnzOracle against baseline models across three dimensions-sequence homology, sequence length, and EC categories-observing that it maintained a leading performance in the vast majority of conditions(Fig. S6-8). While not universally optimal in isolated subsets, this robust overall generalization confirms that the model captures intrinsic biophysical determinants of enzyme stability rather than merely overfitting to sequence memorization.

### Adaptive gating and expert specialization enable robust prediction across biophysical landscapes

To elucidate how the MoE architecture discriminates diverse sequence environments, we visualized the latent feature space projected by the adaptive gating network. The t-SNE embedding (Fig. 4a) reveals that the model spontaneously organizes enzymes into a distinct topological structure driven by task difficulty rather than simple sequence similarity (Fig. S9)^44^. Crucially, this topological separation aligns with the gating network’s routing logic. This discrete routing capability is underpinned by the model’s learning of a continuous physicochemical manifold, indicating that the gating mechanism operates on fundamental biophysical determinants, earned directly from the fused manifold structure, rather than identifying extremophiles as mere statistical anomalies (Fig. S10).

**Fig. 4.**
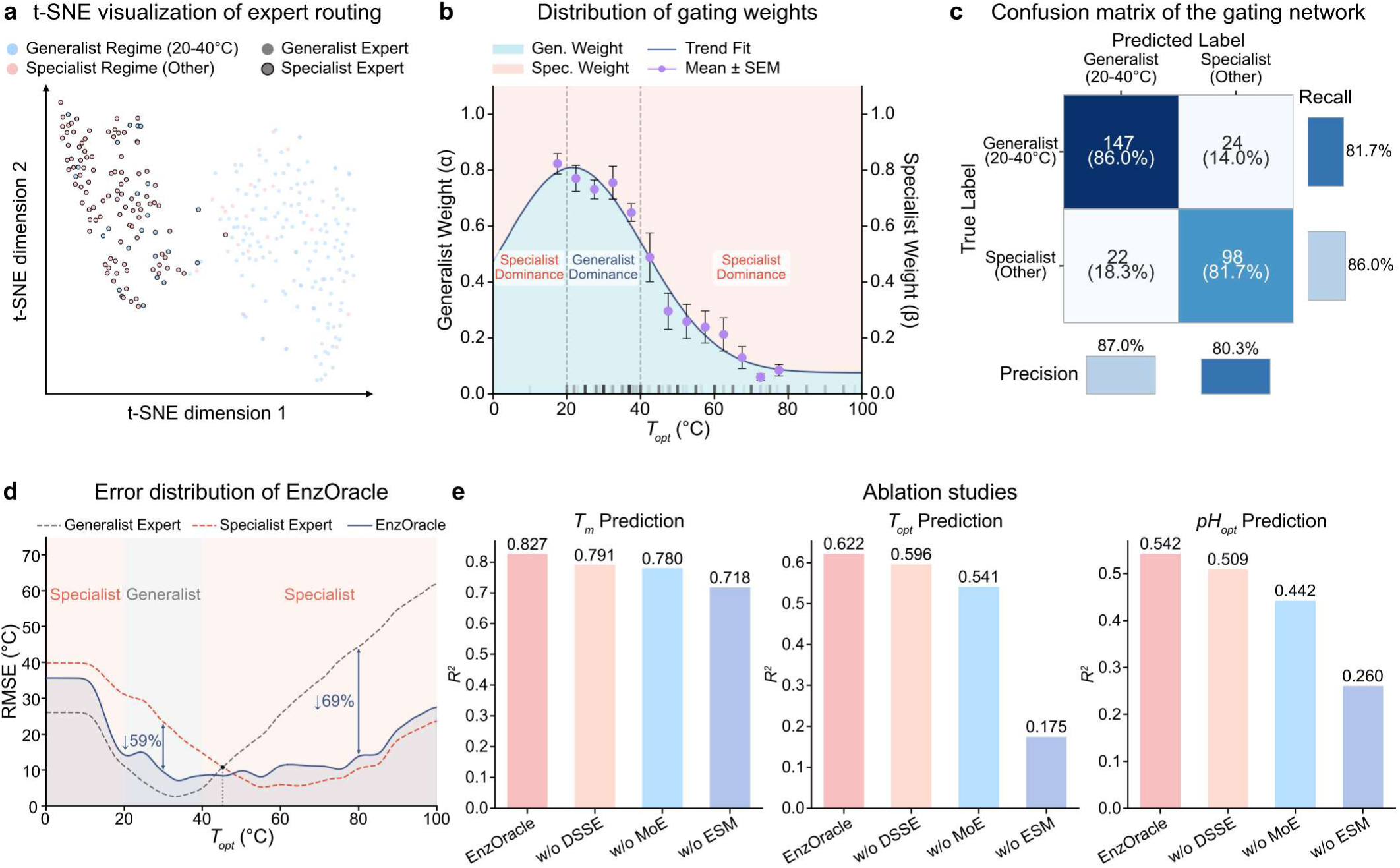
Adaptive gating mechanism and expert specialization. **a**, Latent space visualization of the expert routing logic for *T_opt_*. Blue and pink points represent enzymes belonging to the Generalist Regime and Specialist Regime, respectively. Data points with a black border indicate samples assigned to the Specialist Expert by the gating network. **b**, Gating probability distribution for *T_opt_*. The Generalist Expert weight (*α*, blue trend line) follows a bell-shaped distribution. Purple data points represent the mean gating probability calculated within discrete 5°C bins, with error bars indicating the standard error of the mean (s.e.m.). The background shading visually demarcates the regions of predictive dominance: light blue indicates Generalist Expert dominance (*α*>0.5), while light pink indicates Specialist Expert dominance (*α*<0.5). The rug plot along the x-axis visualizes the data density of the independent test set. **c**, Discriminative accuracy of the adaptive gating network for *T_opt_*. The extended confusion matrix evaluates classification performance, displaying absolute sequence counts and row-normalized percentages for Generalist and Specialist regime assignments in the central cells. Color intensity reflects sample density. Marginal bar charts along the right and bottom axes quantify the Recall and Precision for each class, respectively. **d**, Decoupled expert error profiles for *T_opt_*. Comparative analysis of RMSE trajectories for the Generalist Expert (grey dashed), Specialist Expert (red dashed), and the EnzOracle model (solid blue). Black dots mark equilibrium crossover points, while blue arrows quantify the relative error reduction (“rescue effect”) achieved by the gating mechanism. **e**, Ablation study of architectural components. Vertically stacked bar charts display the *R*^2^ scores for *T_m_* (top), *T_opt_* (middle), and *pH_opt_* (bottom). Colors distinguish the full EnzOracle model (pink) from its ablated variants: without the local encoder (w/o DSSE, orange), without the gating module (w/o MoE, blue), and without the foundation model (w/o ESM, purple).

The core innovation of EnzOracle lies in its ability to dynamically modulate expert collaboration based on input context. We analyzed the gating network’s probability distribution (*α*) to decode this routing logic. For *T_opt_* prediction, the gating weights exhibit a characteristic bell-shaped distribution (Fig. 4b), where the network assigns maximal confidence to the Generalist Expert within the Generalist regime (20-40°C). Conversely, as temperatures deviate into the Specialist regime (<20°C or >40°C), the model dynamically recalibrates its expert weights, executing a continuous shift in predictive authority toward the Specialist Expert. Importantly, this adaptive routing extends beyond thermal optima, demonstrating a clearer functional separation of regimes in stability prediction tasks (Fig. S11). A similar data-driven logic governs *pH_opt_*, where the gating network recapitulates the bell-shaped curve, effectively transferring authority to the Specialist Expert when processing inputs from the corresponding Specialist regime.

The efficacy of the MoE architecture hinges on the gating module’s precision in acting as a routing filter. Evaluated as a binary classifier (Fig. 4c), the gating network demonstrated high-fidelity discrimination. It achieved a recall of 81.7% for the Specialist regime, ensuring that long-tail data points are reliably routed to the specialized network. Concurrently, it maintained an 86.0% recall for the Generalist regime, accurately capturing the dense majority of the data distribution. This discriminative robustness extends across tasks (Fig. S12).

To delineate the complementary contributions of the two sub-networks, we evaluated their decoupled error profiles across the functional spectrum (Fig. 4d). The error profiles reveal a distinct crossover point, demarcating the functional boundaries of the two sub-networks. When operating in the Specialist regime, the gating network executes a “performance rescue” by deploying the Specialist Expert to overcome the Generalist’s underfitting, reducing RMSE by 69% at 80°C. Conversely, within the Generalist regime, dominance dynamically reverts to the Generalist to suppress the Specialist’s overfitting, averting a 59% error penalty at 30°C. Consequently, the global trajectory of EnzOracle effectively forms an optimal error envelope along the minimal error profile of both experts. This complementarity is even more pronounced in *T_m_* prediction, where the dichotomy between stability mechanisms is stark (Fig. S13). Similarly, for *pH_opt_*, the architecture demonstrates bilateral robustness, where the Specialist rescues predictions in both acidic (pH=2, 48% reduction) and alkaline (pH=11, 14% reduction) extremes. Finally, we dissected the contribution of each architectural component through a systematic ablation study (Fig. 4e), revealing a consistent hierarchy of feature importance across all three tasks.

### Biophysical interpretability and structural determinants of functional adaptation

To decode the structural determinants autonomously learned by EnzOracle, we visualized the attention weights of the expert networks projected onto 3D structures. For *T_m_* prediction (Fig. S14), the MoE architecture reveals a functional specialization among expert networks, effectively decoupling surface solubility from core rigidity. The Classification Expert primarily targets surface-exposed polar residues, showing a strong preference for Serine (S) and Tyrosine (Y) to assess folding fidelity. Conversely, the Regression Specialists systematically penetrate the protein surface to lock onto the rigid core. This “stiffness-centric” logic extends to *T_opt_* prediction (Fig. S15), yet with a distinct shift in spatial focus from global scaffolds to active-site preservation. Furthermore, for *pH_opt_* (Fig. S16), the model autonomously discovers the principles of electrostatic microenvironment engineering and captures dynamic gating mechanisms essential for proton exchange.

To rigorously eliminate the potential bias of case-specific, we performed a statistical analysis of the structural attention distributions across all three prediction tasks. For *T_m_* prediction, the global attention profile exhibits a monotonically increasing trend from the protein interior to the exterior (Fig. 5a). While classical folding theory emphasizes the hydrophobic core as the primary driver of basic stability, this result is consistent with the principles of extreme thermal adaptation. The hydrophobic core is a universally conserved prerequisite for folding across all thermal regimes, thus offering low discriminative variance for the model. Instead, EnzOracle implicitly recognizes that hyperthermostability is predominantly achieved through surface optimization, specifically via the formation of extensive solvent-exposed salt-bridge networks and enhanced surface electrostatics. When the prediction target shifts from *T_m_* to *T_opt_*, the model redirects its structural focus, exhibiting a distinct right-skewed, bell-shaped attention distribution(Fig. S17). This topological shift encapsulates the biophysical “stability-activity trade-off.” This task-specific biophysical cognition is further corroborated by the attention landscape for *pH_opt_* prediction.

**Fig. 5.**
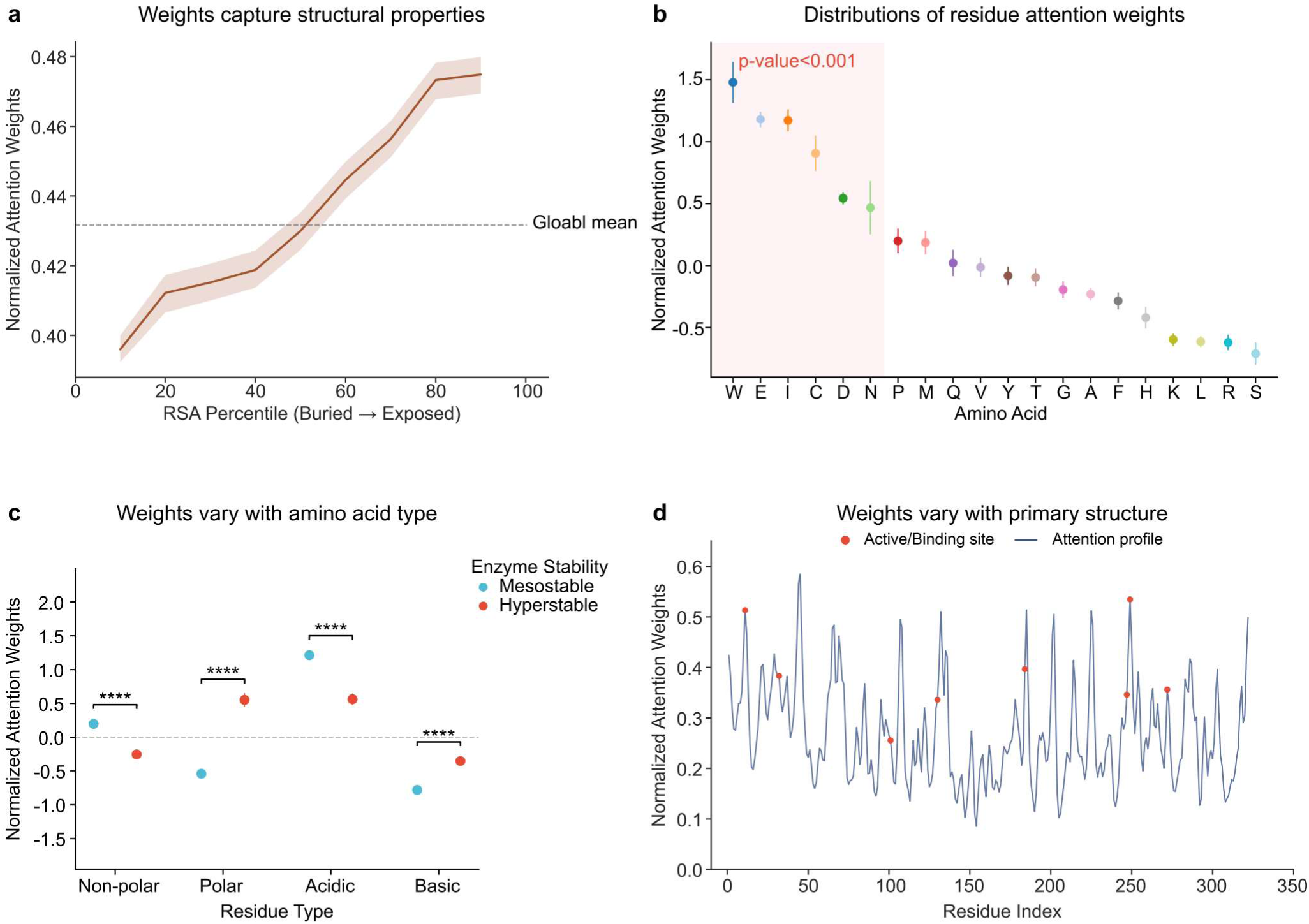
Biophysical interpretability and structural determinants of enzyme stability. **a**, Structural attention landscape for *T_m_*. Gating-weighted normalized attention scores mapped across Relative Solvent Accessibility (RSA) percentiles (0 = deeply buried; 100 = highly exposed)^45^. The solid brown line denotes mean attention weights calculated from n=200 randomly sampled, high-confidence (pLDDT > 70) enzyme structures. This subset maintains a balanced 1:1 ratio of Generalist- and Specialist-dominated samples. The shaded region represents the 95% confidence interval, and the horizontal dashed grey line indicates the global mean. **b**, Global amino acid attention ranking for *T_m_*. Z-score normalized, gating-weighted attention scores across the 20 standard amino acids. Data points denote mean attention weights calculated from the same subset of n=200 enzymes used in Fig. 5a. Error bars indicate the standard error of the mean (s.e.m.). The pink shaded region highlights the top-ranked residues receiving the highest attention (P < 0.001, two-sided Welch’s t-test). **c**, Environment-conditioned attention allocation grouped by residue physicochemical properties for *T_m_*. Normalized attention weights across four residue types. Enzymes are categorized into mesostable (<60°C, blue) and hyperstable (>60°C, red) phenotypes. Data points denote mean attention weights with error bars indicating the standard error of the mean (s.e.m.). Asterisks denote statistical significance across the groups (****P < 0.0001, two-sided Welch’s t-test). **d**, 1D sequence attention mapping for functional hotspots in *T_m_* prediction. Normalized attention weights of residues in the primary structure of an example enzyme (UniProt ID: P61747). Known critical functional loci, including the catalytic proton acceptor (residue 184) and essential cofactor/substrate binding pockets (e.g., residues 11, 101, 130, and 247), are indicated by red dots.

Next, we investigated whether the model’s spatial attention topologically aligns with specific physicochemical properties and observed that EnzOracle develops fundamentally distinct chemical preferences tailored to each predictive task. For *T_m_*, the attention rank is dominated by structural anchors and rigidifying components(Fig. 5b). Conversely, when the prediction target shifts to *T_opt_*, the model markedly shifts attention away from bulky hydrophobic residues. Instead, the highest attention is strictly allocated to polar and catalytic residues(Fig. S18). For *pH_opt_*, the attention landscape is dominated by physiological “pH sensors” possessing titratable side chains (H, C, E), alongside a profound preference for aromatic residues (W, F).

Beyond macroscopic chemical preferences, the MoE system adaptively re-weights features in response to specific extremophilic constraints. We categorized the amino acids into four fundamental physicochemical groups and compared their attention distributions across different environmental phenotypes. For *T_m_* prediction, hyperstable enzymes exhibit a significant upregulation in attention toward Polar and Basic residues compared to their mesostable counterparts(Fig. 5c); conversely, for *T_opt_* prediction thermophilic enzymes exhibit a sharp plummet in attention toward rigidifying Basic residues while maximizing their focus on Polar residues(Fig. S19). Furthermore, the *pH_opt_* expert demonstrates a distinct “electrostatic seesaw” mechanism. It exhibits non-significant variation for non-ionizable residues (Non-polar and Polar), focusing exclusively on electrostatic modulators.

Ultimately, these statistical rationales seamlessly translate into high-resolution micro-level mapping on individual 1D protein sequences(Fig. 5d, Fig. S20). Taken together, these results demonstrate that EnzOracle functions as an interpretable in silico probe, capable of bridging macroscopic biophysical laws with the precise identification of functional and structural determinants at the single-residue level.

### Molecular simulations link EnzOracle attribution to trait-specific adaptation mechanisms

Although attention-based analyses suggested that EnzOracle prioritizes residues with distinct structural and chemical roles across Tm, Topt and pHopt prediction, attention alone cannot establish whether these residues participate in physically meaningful molecular mechanisms. We therefore performed trait-resolved molecular simulations as an orthogonal validation layer. Rather than using MD simulations as an additional benchmark of predictive accuracy, we used them to test whether sequence-derived attribution patterns could be mapped onto independent atomistic observables, including conformational rigidity, catalytic preorganization, hydrogen-bond and salt-bridge persistence, metal coordination, and solvent organization. This design allowed us to interrogate whether EnzOracle captures transferable biophysical principles underlying enzyme environmental adaptation.

The PETase analyses show that EnzOracle attention hotspots coincide with residues and structural regions involved in preserving productive substrate-binding geometries, oxyanion-hole stability, and transition-state stabilization. These findings support the view that, for *T_opt_* prediction, EnzOracle captures local dynamic constraints required for catalysis under thermal stress(Fig. 6, S21).

**Fig. 6.**
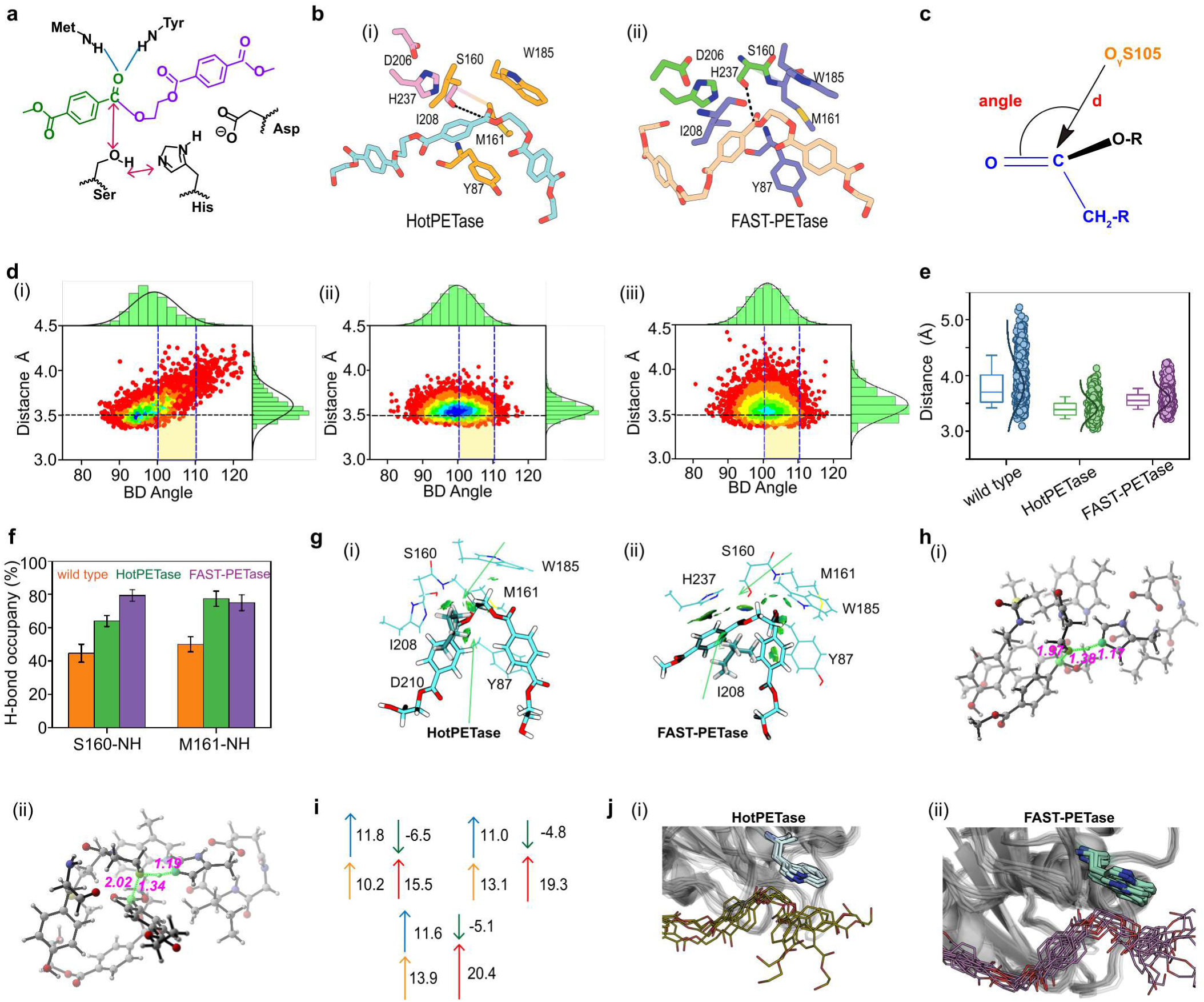
Structural and conformational analysis of PET hydrolases based on molecular dynamics simulations. **a**. Proposed catalytic mechanism of PET hydrolysis, highlighting the acylation step initiated by nucleophilic attack of the catalytic Ser-Oγ atom on the scissile ester carbonyl carbon and stabilization of the tetrahedral intermediate by the oxyanion hole. **b**, Representative MD-derived conformations of PET-bound (**i**) FAST-PETase, and (**ii**) HotPETase, with key functional groups involved in ester bond cleavage and substrate recognition highlighted. **c**, Definition of the prereaction-reactive state (PRS), including the nucleophilic attack distance d(C···Oγ) and the corresponding Bürgi-Dunitz approach angle. **d**, Distributions of d(C···Oγ) for wild-type *Is*PETase, FAST-PETase, and HotPETase across MD trajectories. FAST-PETase and HotPETase maintain narrower near-attack distributions than wild-type *Is*PETase. **e**,**f**, hydrogen-bonding analysis of the oxyanion hole. Donor-acceptor distance distributions and occupancy profiles quantify the stability of catalytic hydrogen-bond networks during MD simulations. **g**, IGMH analysis of noncovalent interactions between PET and the enzyme active-site pocket, highlighting stabilizing contacts around the scissile ester bond. **h**, Optimized transition structures for the acylation step in HotPETase and FAST-PETase. Key interatomic distances are shown in Å. **i**, Distortion/interaction activation strain analysis based on theozyme DFT cluster models, decomposing the acylation barrier into distortion and interaction components. **j**, analysis of W185 conformational dynamics and structural superposition of MD-derived conformational ensembles, illustrating differences in active-site plasticity and substrate-recognition geometry among PETase variants.

To validate determinants associated with melting temperature, we selected Thermus thermophilus UV damage endonuclease (UVDE) as a representative thermostable enzyme system(Fig. S22, S23). Model attribution identified N107 as a high-importance residue located within a densely packed internal region of the UVDE scaffold. To evaluate its structural role, we constructed an N107A variant in silico and compared its dynamics with the wild-type enzyme using atomistic MD simulations based on the crystal structure of UVDE. Structural analysis revealed that N107 acts as a rigidity-anchoring residue within the UVDE scaffold.

To evaluate whether EnzOracle captures determinants of pH adaptation, we performed MD simulations on an alkaline pectate lyase under protonation regimes corresponding to pH 7.0 and pH 10.0(Fig. S22). Results indicate that pH adaptation arises primarily from redistribution of electrostatic interactions rather than changes in protonation states alone. EnzOracle-prioritized residues form persistent interaction networks that buffer electrostatic perturbations, mapping sequence-derived importance to pH-dependent structural behavior.

Collectively, these analyses demonstrate that EnzOracle-derived attention patterns correspond to physically meaningful structural and dynamical features. Across all three systems, residues assigned high importance by the model map onto regions governing catalytic flexibility, global stability, or electrostatic adaptation. This correspondence suggests that EnzOracle captures transferable biophysical principles underlying enzyme adaptation rather than relying solely on dataset-specific statistical correlations. Conversely, the framework provides a means to identify cases where model attribution does not align with physical behavior, thereby defining the limits of interpretability. Taken together, the “1+2” validation strategy establishes MD as a critical evidential layer for assessing the mechanistic validity of sequence-based enzyme property prediction, linking model attribution to experimentally interpreable molecular mechanisms across diverse environmental regimes.

## Discussion

Biocatalyst engineering has long been bottlenecked by the intrinsic dissonance between natural evolutionary selection and industrial operational requirements. Monolithic deep learning models, while powerful, inevitably fall victim to the “regression to the mean” trap when faced with this severe data asymmetry, often treating rare extremophilic features as statistical noise. EnzOracle effectively addresses this paradigm through a CG-MoE architecture. EnzOracle achieves high predictive accuracy across both the core data distribution and challenging long-tail extremes, without relying on external data.

The necessity of this decoupled architecture stems from the fundamental biological reality that mechanisms conferring canonical functioning and extremophilic resilience are often physically mutually exclusive. In monolithic architectures, optimizing for these opposing traits simultaneously induces destructive gradient interference. The adaptive gating module in EnzOracle acts as a critical semantic filter. By routing sequences based on an autonomously learned physicochemical manifold, the gating mechanism orchestrates a dynamic “performance rescue.” This strategy not only shields the Generalist Expert from outlier interference but also prevents the Specialist Expert from severe extrapolation errors, enabling smooth and accurate interpolation across the entire fitness landscape.

Beyond predictive fidelity, EnzOracle illuminates the “black-box” nature of deep sequence matching by demonstrating the emergence of task-specific biophysical first principles. The model’s attention mechanism autonomously discovers the classic “stability-activity trade-off”: for *T_m_* prediction, it prioritizes rigidifying surface salt bridges and hydrophobic anchors, whereas for *T_opt_*, it shifts focus away from these bulky residues in favor of the dynamic polar networks essential for catalytic turnover. Notably, for *pH_opt_*, the model exhibits a distinct “electrostatic seesaw,” dynamically inverting its focus on acidic and basic residues in response to environmental proton availability. The precise alignment of these macroscopic statistical trends with micro-level functional hotspots confirms that EnzOracle extracts true physicochemical determinants rather than spurious homological artifacts.

This robust physicochemical cognition directly translates into tangible industrial value. In the context of virtual screening, EnzOracle’s ability to maintain high screening success rates under strict industrial tolerances significantly reduces the experimental validation burden associated with metagenomic mining. Furthermore, the high-resolution structural attention maps generated by the model can function as an in silico probe. By pinpointing the specific residues that dictate extremophilic adaptation, EnzOracle provides actionable blueprints for the rational design and directed evolution of next-generation biocatalysts.

The molecular simulation analyses further strengthen the mechanistic interpretation of EnzOracle. Rather than functioning as post hoc visualization, the simulations provided an independent physical test of whether model attribution corresponds to measurable molecular behavior. The agreement between attention hotspots and trait-specific observables (core rigidity for Tm, catalytic preorganization for *T_opt_*, and electrostatic/hydration remodeling for *pH_opt_*) suggests that the model’s internal representations are aligned with physically meaningful determinants of enzyme adaptation. This is particularly important for enzyme engineering, where interpretability is valuable only when it can nominate residues whose structural or dynamic roles are mechanistically plausible.

Despite these advances, we acknowledge current limitations in sequence-based predictive paradigms. While foundational models like ESM-2 implicitly encode structural information, enzymatic function is fundamentally governed by dynamic 3D conformational changes, explicit solvent interactions, and the presence of essential cofactors-factors that are currently only inferred indirectly. Additionally, while *T_opt_* and *pH_opt_* are critical operational parameters, comprehensive industrial scale-up requires the precise prediction of complex kinetic behaviors, including turnover rates (*k_cat_*), binding affinities (*K_m_*), and product inhibition constants (*K_i_*). To bridge this gap between static sequence prediction and dynamic catalytic modeling, physical simulations are often employed; however, the present MD and DFT analyses should be interpreted as representative physical validations rather than exhaustive causal proofs across all enzyme families. Such simulations depend heavily on force-field accuracy, protonation-state assignments, and sampling length, while cluster models inevitably simplify the full enzymatic environment. Consequently, future iterations of the EnzOracle framework will integrate geometric deep learning over explicit 3D structures with QM/MM free-energy simulations and large-scale mutational assays. This synergy will evolve the present attribution-validation framework beyond stability and optima prediction, ultimately forging a closed-loop, generative platform for mechanism-guided enzyme design.

In summary, EnzOracle establishes a distribution-aware framework for predicting enzyme physicochemical parameters directly from sequence while simultaneously enabling mechanistic interpretation of environmental adaptation. Through an orthogonal molecular dynamics validation strategy spanning catalytic temperature optimization, thermal stability, and alkaline adaptation, we demonstrate that residues prioritized by the model correspond to dynamically constrained structural cores, preorganized catalytic conformational ensembles, and pH-dependent electrostatic interaction networks, respectively.

## Methods

### Data preparation

#### Data sources and sequence curation

To ensure rigorous benchmarking and prevent data leakage, all training and evaluation datasets were strictly sourced from established official benchmarks. For *T_m_* and *T_opt_* predictions, datasets were obtained from the DeepTM^27^ and TOMER^24^ benchmarks, respectively. For *pH_opt_* prediction, data was retrieved from the EpHod^23^ database.

#### Stratified splitting strategy

To evaluate the model’s generalization capabilities impartially, the original independent test sets from the respective benchmarks were preserved completely intact (n=1,550 for *T_m_*, n=291 for *T_opt_*, and n=1,971 for *pH_opt_*). For the *pH_opt_* task, we maintained the pre-defined splits from EpHod, resulting in 7,124 training and 760 validation sequences.For the *T_m_* and *T_opt_* tasks, which lacked official validation partitions, we developed a robust stratified binning strategy to extract a 10% validation set from the training corpus. Specifically, the data was discretized into fixed-width bins (5°C intervals for *T_m_* and 10°C intervals for *T_opt_*) to ensure sufficient sampling across the entire functional landscape. This stratified sampling yielded final splits of 5,616 training / 624 validation sequences for *T_m_*, and 2,363 training / 263 validation sequences for *T_opt_*. Statistical equivalence between the training, validation, and testing distributions was subsequently confirmed via the Kolmogorov-Smirnov (KS) test and Kullback-Leibler (KL) divergence evaluations.

#### Regime definition and expert labeling

A core prerequisite for the CG-MoE framework is the explicit categorization of sequences into the data-dense Generalist regime versus the data-sparse Specialist regime. To train the AGN and route inputs to the appropriate regression expert, binary labels were programmatically assigned based on the empirical data distribution. Sequences falling within the data-abundant core bounds (specifically 50≤*T_m_*≤ 60°C, 20≤*T_opt_*≤40°C, and 7≤*pH_opt_*≤8) were labeled as “1” and assigned to the Generalist Expert. Conversely, sequences falling outside these boundaries, representing the sparse distributional tails, were labeled as “0” and allocated to the Specialist Expert. This discrete labeling strategy ensures that the specialized sub-network is optimized exclusively for long-tail edge cases without gradient interference from the data-dense majority.

### Architecture of the EnzOracle Framework

The EnzOracle framework was designed to independently capture structural rules from the core data distribution and its long-tail edge cases. To prevent gradient interference during backpropagation, the framework instantiates three completely decoupled neural architectures for each prediction task: the AGN, the Generalist Expert (α), and the Specialist Expert (β). While these three sub-networks function independently, they share a unified architectural topology termed the Hybrid Global-Local Representation Engine (HGL-RE), which processes evolutionary and sequential modalities in parallel. Specific architectural hyperparameters (e.g., layer depths, hidden dimensions, and attention heads) were individually optimized for each sub-network and are detailed in Supplementary Table 1.

#### Pre-trained Protein Language Model

To infuse deep evolutionary semantics into the representation, we employed the ESM-2 foundational protein language model (the esm2_t33_650M_UR50D variant)^33^. The ESM-2 model was kept strictly frozen during training. For an input amino acid sequence of length *L*, the contextual embeddings were extracted from the terminal 33rd layer, yielding a high-dimensional evolutionary feature matrix *E ∈ R^L*1280^*, where the embedding dimension (*d_esm_*) is 1280.

#### Deep Semantic Sequence Encoder

To extract high-dimensional latent representations from discrete amino acid sequences, all nine constituent models across the three predictive tasks (*T_m_*, *T_opt_*, and *pH_opt_*) share a universally parameterized semantic sequence encoder. Initially, the raw sequence is mapped into a continuous latent space utilizing a learnable embedding layer with a dimension of *d_model_*=128. To inject relative position information, standard sinusoidal positional encodings are superimposed onto the embeddings, followed by a dropout layer (*p* = 0.2) for regularization^42^.

The position-aware sequence embeddings are subsequently processed through a stack of *N*=3 consecutive Transformer encoder layers^42^. Each layer fundamentally comprises a Multi-Head Self-Attention (MHSA) module and a Position-wise Feed-Forward Network (FFN). For the MHSA module, we implemented *h*=4 independent attention heads. The input representations are linearly projected into queries (*Q*), keys (*K*), and values (*V*), where the dimensionality for each head is strictly set to *d_k_*=*d_v_*=64. To ensure that the self-attention mechanism strictly ignores the zero-padded regions of variable-length sequences, a dynamic padding mask (*M*) is applied prior to the softmax normalization. The scaled dot-product attention for each head is mathematically formulated as:

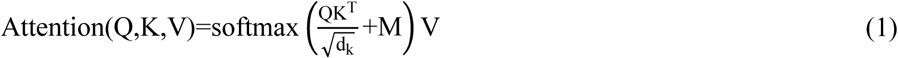

where *M_ij_*=-∞ if the *j*-th token is a padding token, and 0 otherwise. The outputs from all *h* heads are concatenated and linearly projected back to the original dimension *d_model_*.

Following the MHSA, the representation undergoes transformation via the FFN, which utilizes an inner hidden dimension of *d_ff_*=512. The FFN executes two linear transformations devoid of bias, separated by a ReLU activation and a dropout layer (*p*=0.1). Crucially, to facilitate stable gradient propagation in deep architectures, both the MHSA and FFN sub-layers are encapsulated by residual connections and Layer Normalization^46,47^. The finalized output of this encoder matrix directly serves as the input query sequence (*H_seq_*) for the downstream Evolutionary-Contextual Fusion Interface (ECFI).

To enrich the sequence semantics with structural priors without requiring explicit alignments, the learned representations are passed into the Evolutionary-Contextual Fusion Interface (ECFI). This module executes a cross-attention mechanism wherein the sequence embeddings act as the Query (*Q*), while the dense evolutionary representations derived from the frozen ESM-2 act as the Key (*K*) and Value (*V*). The fused multi-modal representation is then branched into two complementary feature extraction pathways:

1. **Local Pathway (1D-CNN)**: To capture spatially localized functional motifs, the fused embeddings are processed through parallel 1-dimensional convolutional layers (1D-CNN). We deployed multiple kernel sizes (*k* ∈(3, 5, 7)) to ensure multi-scale receptive fields, followed by LeakyReLU activation and max-pooling over time.
2. **Global Pathway (Attention Pooling):** Concurrently, the sequence is fed through additional Transformer layers followed by a self-attention pooling mechanism, dynamically aggregating long-range stability constraints into a fixed-length global vector. The outputs from both the global and local pathways are concatenated to form the definitive comprehensive enzyme fingerprint.

#### Adaptive Gating Network and Dual-Expert Routing

The routing logic of the MoE system is entirely governed by the gating module^48^. The enzyme fingerprint extracted by the HGL-RE of the gating module is fed into a classification head with a Sigmoid activation, outputting a continuous scalar probability (*W_prob_* ∈ [0, 1]). This scalar quantitatively represents the likelihood of the query sequence belonging to the Generalist regime.

Simultaneously, the Generalist Expert (*α*) and the Specialist Expert (*β*) independently process the identical sequence through their respective, fully decoupled HGL-RE networks to produce distinct scalar predictions (*Ŷ_Generalist_* and *Ŷ_Specialist_*). The ultimate functional parameter (*Ŷ_final_*) is calculated via a dynamic, probability-weighted summation (late fusion):

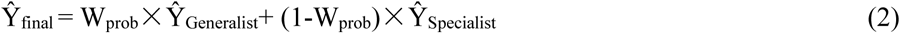

This conditional inference strategy ensures smooth predictive interpolation across the biophysical landscape while strictly maintaining the semantic purity of the individual expert manifolds.

### Training routine and task-specific optimization

Neural network models were constructed utilizing the PyTorch v1.12.1 framework. Training was executed on NVIDIA RTX 3090. To strictly prevent gradient interference between Generalist regime and Specialist regime feature learning, all nine sub-networks across the three property prediction tasks (*T_m_*, *T_opt_*, *pH_opt_*) were trained completely independently from scratch.

Crucially, because the biophysical landscapes and data distributions vary fundamentally across different environmental regimes, we did not employ a monolithic training protocol. Instead, optimization algorithms, loss formulations, batching strategies, and learning rate schedules were highly heterogeneous and uniquely tailored to the specific convergence requirements of each individual model. A comprehensive breakdown of the precise hyperparameters for all nine sub-networks is provided in Supplementary Table 2. Gradient estimation strategies and learning rate schedules also varied by model capacity and task complexity. Where VRAM constraints and gradient stability necessitated it, an effective batch size was simulated utilizing a physical mini-batch size coupled with 4 gradient accumulation steps. To mitigate overfitting, early stopping was universally enforced across all training routines, with the patience threshold optimized per task.

### Performance evaluation

#### Quantitative metrics

To systematically quantify the predictive fidelity and generalization capabilities of the continuous regression experts, we employed four standard statistical criteria: the, RMSE, MAE, and the Pearson correlation coefficient (Pearson *r*). These metrics collectively assess the variance explanation, average prediction deviation, and linear correlation between the experimental ground truths and the computational predictions.

For the AGN, which functions as the architectural gating mechanism via binary classification, the discriminative performance was evaluated using Recall (Sensitivity) and Precision (Positive Predictive Value, PPV). These classification metrics are mathematically formulated as:

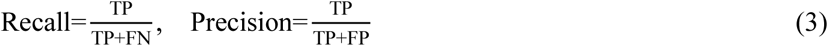

where *TP*, *FN*, and *FP* denote the cumulative counts of true positives, false negatives, and false positives across the test distribution, respectively.

#### Regression Error Characteristic (REC) analysis

Standard regression metrics (e.g., RMSE) are highly sensitive to extreme outliers and often fail to reflect a model’s practical utility in industrial biocatalyst design. To address this, we conducted a REC analysis. The REC curve plots the absolute error tolerance on the x-axis against the cumulative percentage of predictions falling within that tolerance on the y-axis, providing a comprehensive visualization of the model’s error cumulative distribution function.

From the REC curves, we explicitly defined the “Screening Success Rate,” representing the proportion of sequence predictions where the absolute error |*ŷ_i_*−*y_i_*| is strictly less than or equal to a predefined industrial tolerance threshold (*τ*). To reflect authentic reactor operation tolerances and directed evolution constraints, *τ* was rigorously established at ±5°C for thermal parameters (*T_m_* and *T_opt_*) and ±0.5 units for *pH_opt_*. This targeted metric directly translates computational accuracy into tangible virtual screening reliability.

#### Baseline comparisons

To rigorously establish the SOTA standing of the EnzOracle framework, comprehensive head-to-head benchmarking was performed against twelve recently published computational models. The baseline competitors included DeepSTABp^25^, TemStaPro^26^, ProTstab2^21^, and DeepTM^27^ for *T_m_* prediction; Seq2Topt^30^, TOMER^24^, DeepET^28^, and Preoptem^29^ for *T_opt_* prediction; and OphPred^32^, Catopt^31^, EpHod^23^, and Seq2pHopt^30^ for *pH_opt_* prediction. To preclude any potential data leakage and ensure an absolutely impartial, head-to-head comparison, all baseline predictions were independently regenerated on our identical, fully held-out test sets utilizing their officially released open-source repositories or publicly accessible web servers.

### Interpretability analysis and structural mapping

#### Attention extraction and normalization

To elucidate the biophysical first principles autonomously learned by the EnzOracle framework, we extracted the raw attention weights directly from the attention pooling module (att_pool) within the ECFI. To ensure the interpretability of these learned weights across different analytical contexts, two distinct normalization strategies were employed. For the single-sequence 3D structural mapping, the raw attention scores were scaled using Min-Max normalization to bound the values strictly between 0 and 1, providing a clear visual gradient of functional hotspots. Conversely, for the global statistical analysis conducted across a representative cohort of 200 diverse enzymes, a within-sequence Z-score standardization was applied. This critical step eliminates the inherent attention-dilution bias introduced by varying sequence lengths, allowing for a fair, sequence-independent comparison of residue-level importance across different enzyme families.

#### 3D structural analysis and solvent accessibility

To bridge the 1D sequence attention with 3D spatial conformations, high-quality protein structures were systematically sourced. We prioritized experimentally resolved structures from the Protein Data Bank (PDB)^49^. In the absence of experimental data, computational structures were retrieved from the AlphaFold Protein Structure Database (https://alphafold.ebi.ac.uk/), and for remaining sequences, structures were generated de novo using ESMFold^33,50^. To ensure spatial reliability, all computationally predicted structures were strictly filtered using a high-confidence threshold (average pLDDT>70).

To quantify the spatial microenvironment of highlighted residues, the absolute Solvent Accessible Surface Area (SASA) was computed using the DSSP algorithm (specifically, the mkdssp executable, version 2.0.4). The RSA was subsequently derived by dividing the absolute SASA by the theoretical maximum SASA specific to each of the 20 standard amino acids. The final calculated RSA values were mathematically clipped to a continuous range of [0, 1]. All high-resolution mapping of attention weights onto 3D protein topologies was visualized and rendered using PyMOL (Version 3.0.0, Schrödinger, LLC).

#### Statistical tests

To rigorously validate the observed macroscopic biophysical trends, all comparative analyses for statistical significance were conducted using the two-sided Welch’s t-test. This method was specifically selected as it does not assume equal population variances, thereby providing a more robust evaluation of the highly skewed extremophilic distributions.

### MD simulations

MD simulations were performed using the Amber 22 program suite, employing the ff19SB and GAFF force fields for protein and ligand modeling, respectively^51,52^. Protonation states of titratable residues (His, Glu, Asp) were determined using PROPKA via the PDB2PQR server^53^. We selected three PET-degrading enzymes: wild-type *Is*PETase, FAST-PETase, and HotPETase. FAST-PETase is a machine-learning-guided variant carrying five substitutions relative to IsPETase (S121E, D186H, R224Q, N233K, and R280A)^35^. HotPETase, is a highly thermostable IsPETase variant with a reported Tm of 82.5°C and excellent PET-hydrolytic activity^54^. The initial protein structures for the enzymes used in this study, including *Is*PETase (PDB ID: 5XH3), HotPETase (PDB ID: 7QVH), and FAST-PETase (PDB ID: 7SH6), were obtained from previously reported X-ray crystal structures. The PET substrate model was a PET trimer (2-hydroxyethyl-(mono-hydroxyethyl terephthalate)₃), which aligns with previous simulation studies^55^. To ensure robust sampling and reproducibility, three independent 500 ns production trajectories were generated for each enzyme system^56^. Trajectory analyses were performed using CPPTRAJ^57^. Non-covalent interactions were further analyzed using the Independent Gradient Model based on Hirshfeld Partition (IGMH) methods^58^. Visualization and map generation were conducted using VMD 1.9.3 and Multiwfn _3.758,59._

## Data availability

All datasets generated and analyzed in this study are publicly available at https://github.com/kiki77-gq/EnzOracle.

## Code availability

The custom source code used to train and evaluate the EnzOracle models is freely accessible at https://github.com/kiki77-gq/EnzOracle.

## Supporting Information

Supporting information is available for this paper.

## Acknowledgements

The authors gratefully acknowledge the members of the Wei Lab for their insightful discussions, technical assistance, and continuous support throughout the course of this study.

## Author Contributions

Qi Gao: conceptualization, study design, algorithm development, experimentation, data analysis, writing original draft. Zhongcheng Fang: methodology design, data curation and software implementation, Yajing Yuan: software implementation. Mengyuan Jin: software implementation. Heqi Sun: software implementation. Zhennan Peng: software implementation. Lan Yang: software implementation. Jiayi Li: study design, methodology design, writing-review and editing. Dong-Qing Wei: : supervision, resources, writing-review, and supervision.

