## Supplementary Information for "EnzOracle: Mechanism-aware prediction of enzyme environmental adaptation via a classification-guided mixture-of-experts framework"

### Supporting Information

#### Table of Contents

##### Supplementary Figures

1 The misalignment between evolutionary selection and industrial requirements
2 Distributional consistency between training and evaluation sets
3 Residual analysis and systematic bias assessment
4 Functional generalization across enzyme families
5 REC curves for extremophilic variant screening
6 Generalization capability across sequence identity thresholds
7 Robustness against sequence length variations
8 Broad applicability across Enzyme Commission (EC) categories
9 Latent space routing logic for stability and pH tasks
10 Emergence of continuous physicochemical manifolds
11 Adaptive gating dynamics for $T_m$ and $pH_{opt}$
12 Discriminative reliability for $T_m$ and $pH_{opt}$
13 Complementary expert error profiles for $T_m$ and $pH_{opt}$
14 Visualization of structural attention weights for $T_m$ prediction
15 Visualization of structural attention weights for $T_{opt}$ prediction
16 Visualization of structural attention weights for $pH_{opt}$ prediction
17 Structural attention landscapes for $T_{opt}$ and $pH_{opt}$
18 Global amino acid attention rankings for $T_{opt}$ and $pH_{opt}$
19 Environment-conditioned attention allocation grouped by residue physicochemical properties
20 1D sequence attention mapping for functional hotspots
21 High-temperature RMSD analysis (350 K)
22 MD validation of melting-temperature and pH-adaptation determinants in UVDE and PelA
23 Analysis of the distance change between residues and $Ca^{2+}$ centers during MD simulation.

##### Supplementary Tables

1 Task-specific architectural hyperparameters for the decoupled expert networks
2 Task-specific training hyperparameters and optimization objectives for the decoupled expert networks

**Supplementary Note：Formulation of Objective Functions**

**Supplementary Note: MD simulations**

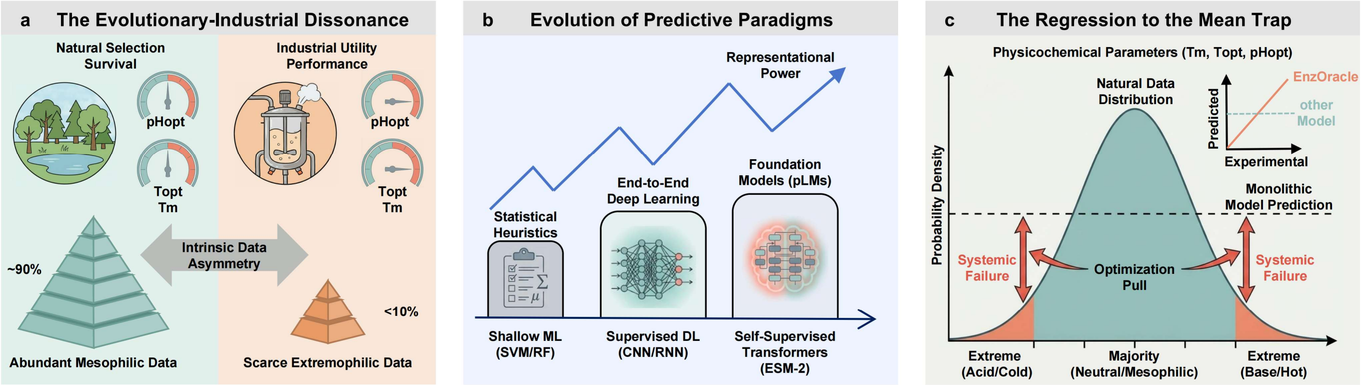


**Supplementary Fig. 1 | The misalignment between evolutionary selection and industrial requirements.**

**a**, The intrinsic misalignment between natural evolution and industrial necessity. Natural selection favors fitness under physiological conditions (mesophilic, neutral pH), resulting in a long-tail distribution where high-value extremophilic enzymes are statistically scarce compared to the data-dense canonical majority. **b**, The transition of predictive methodologies. **c**, The "regression to the mean" phenomenon.

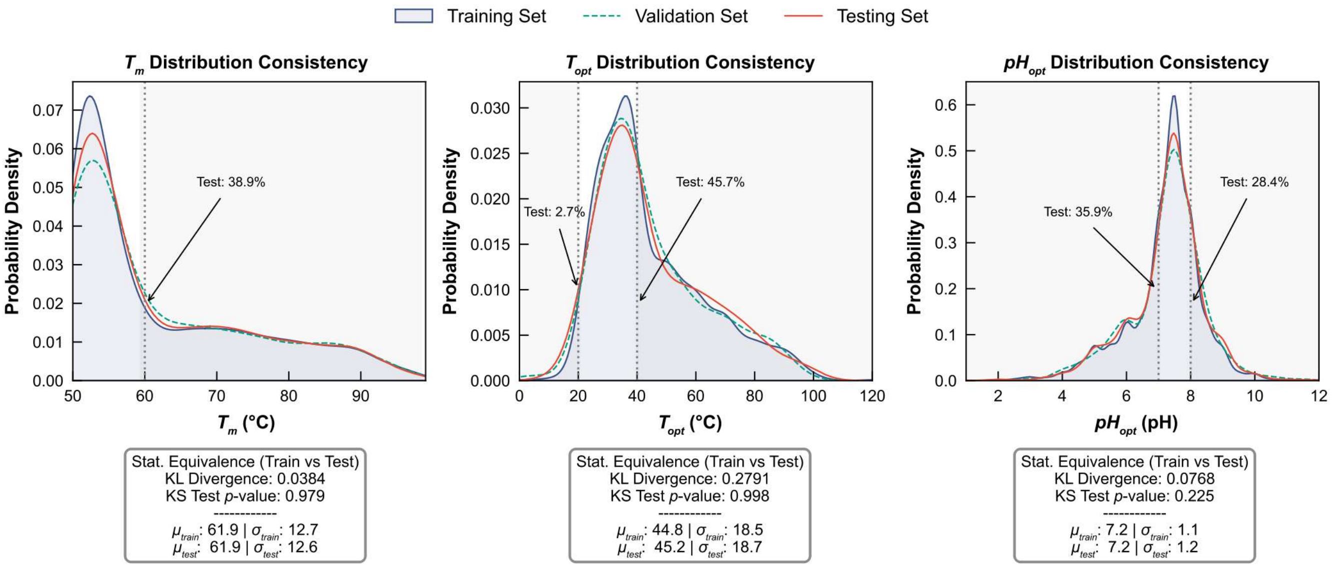


**Supplementary Fig. 2 | Distributional consistency between training and evaluation sets.**
Kernel density estimation (KDE) plots comparing the probability distributions of training (grey shaded area), validation (dashed green line), and testing (solid red line) datasets for $T_m$ (left), $T_{opt}$ (middle), and $pH_{opt}$ (right). Data splitting was performed via stratified sampling to ensure that the evaluation sets statistically mirror the underlying distribution of the training corpus. Statistical equivalence was quantified using Kullback-Leibler (KL) divergence and the Kolmogorov-Smirnov (KS) test (results annotated in insets). High $p$-values ($p > 0.05$) and low divergence scores confirm no significant distributional shift. Arrows highlight the explicit coverage of extremophilic tail regions in the independent test set, ensuring that the reported metrics reflect performance in data-scarce regimes.

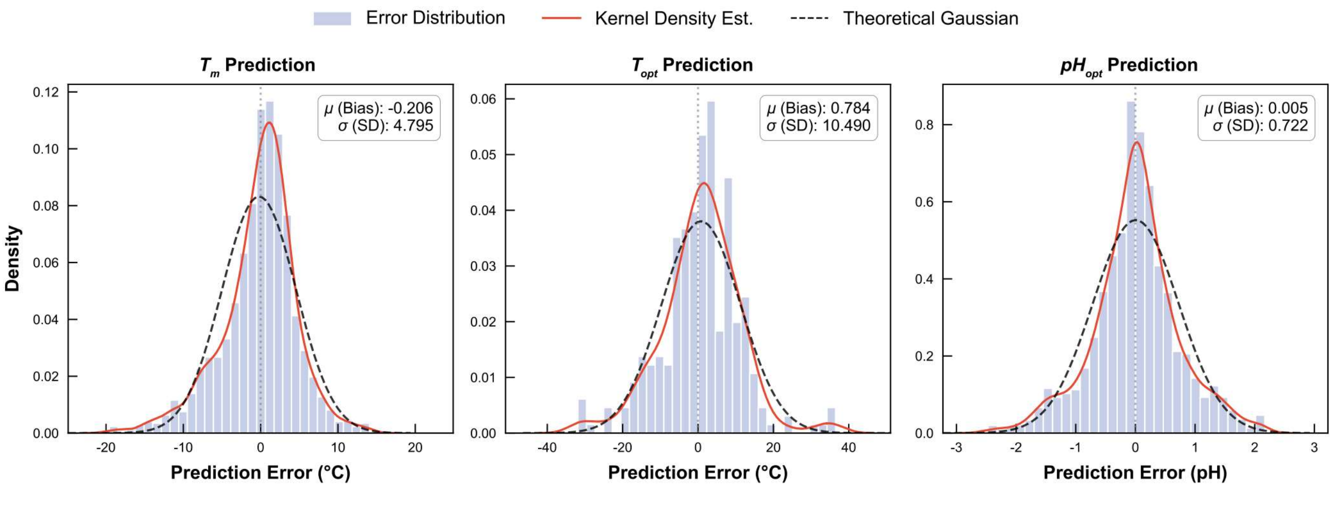


**Supplementary Fig. 3 | Residual analysis and systematic bias assessment.**
Error distribution histograms for $T_m$ (left), $T_{opt}$ (center), and $pH_{opt}$ (right) predictions on the independent test set. The red solid line represents the empirical kernel density estimate, while the black dashed line represents a theoretical Gaussian fit derived from the residual statistics ($\mu$, $\sigma$). Statistical insets display the mean bias ($\mu$) and standard deviation ($\sigma$).

While the consistent alignment of median errors along the zero line (dashed), the variance in prediction error exhibited distinct task-specific patterns. Specifically, $T_{opt}$ predictions showed higher dispersion for Isomerases (EC 5), likely reflecting the extreme structural heterogeneity inherent to this class, whereas $pH_{opt}$ predictions maintained consistent tightness across all functional categories. Of particular significance is the model's performance on Translocases (EC 7). Despite this class being predominantly composed of membrane-associated proteins with distinct hydrophobic profiles and scarce experimental data, EnzOracle maintained high predictive fidelity. This functional robustness implies that the model has successfully decoupled the learning of universal thermodynamic stability rules (e.g., packing density, electrostatic networks) from local catalytic mechanisms, preventing overfitting to specific active site geometries associated with particular reaction types.

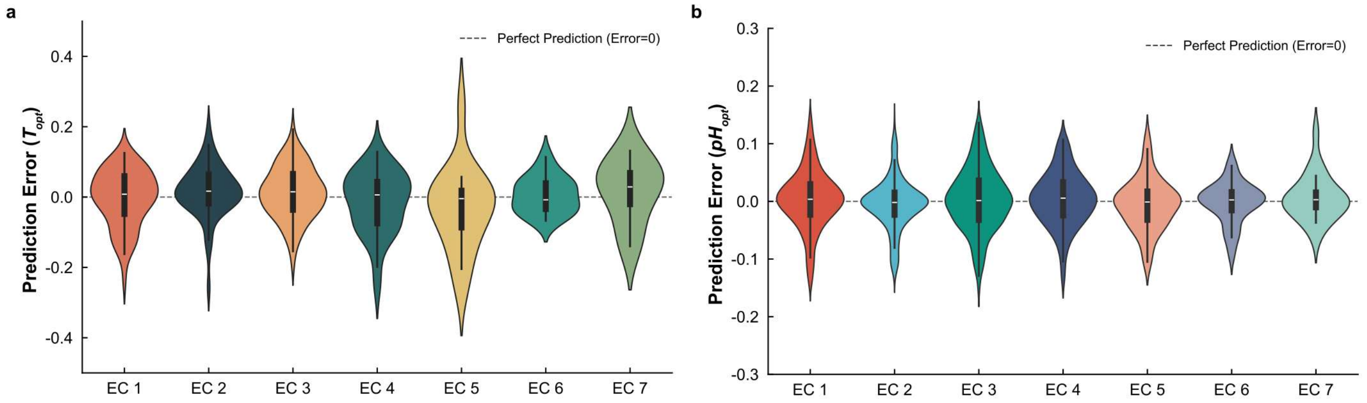


**Supplementary Fig. 4 | Functional generalization across enzyme families.**

Violin plots illustrating the distribution of range-normalized prediction errors for **a**, $T_{opt}$ and **b**, $pH_{opt}$, stratified by Enzyme Commission (EC) main classes (EC 1-7). To facilitate comparative visualization across scales, raw errors were normalized by the effective physiological range (120°C for $T_{opt}$ and 14 units for $pH_{opt}$). Furthermore, to prevent visual distortion by extreme outliers, the distributions were truncated to the 1st-98th percentiles for $T_{opt}$ and the 1st-99th percentiles for $pH_{opt}$. Within each distribution, the central white line denotes the median, the thick vertical bar spans the interquartile range (IQR), the thin vertical line extends to the 1.5× IQR bounds (whiskers), and the symmetric outer contour represents the kernel density estimation of the prediction residuals.

In the identification of hyperstable variants, EnzOracle achieved a success rate of 53.5% within the industrial tolerance of ±5°C, delivering a 6.6% relative improvement over the SOTA. Similarly, for variants adapted to extreme pH environments, EnzOracle maintained a success rate of 41.8% within a tight margin of ±0.5 pH units, surpassing the baseline by approximately 7.5%.

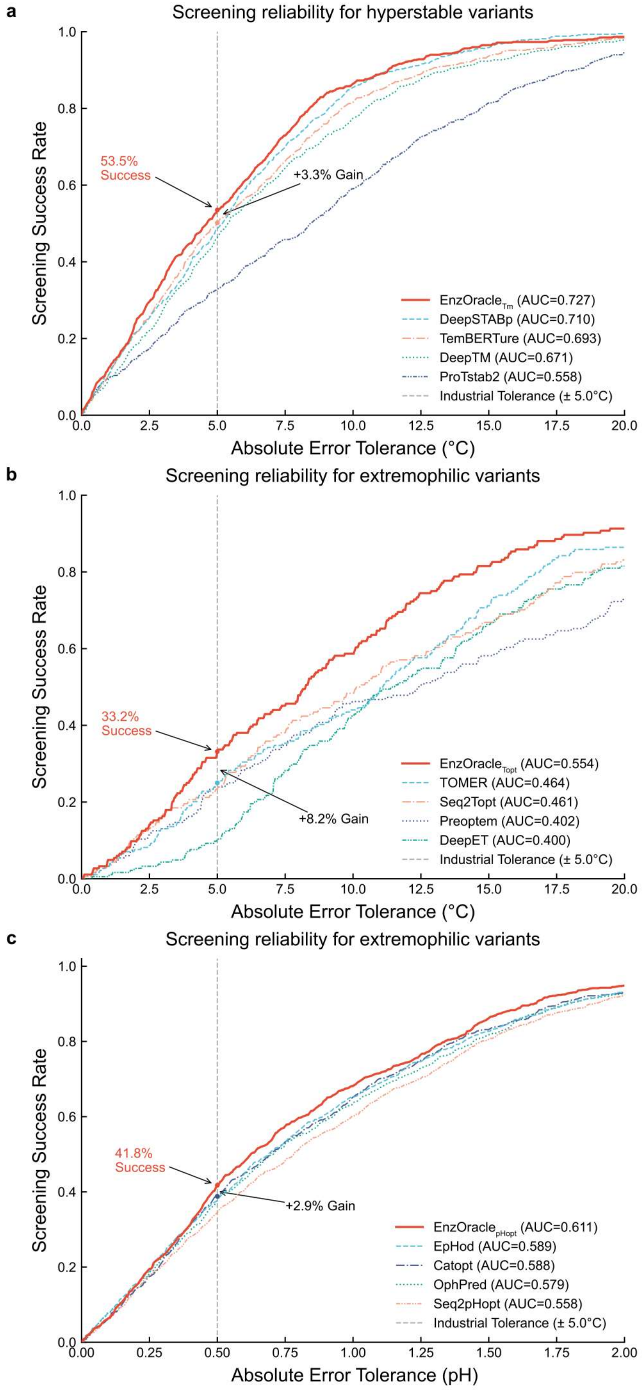


**Supplementary Fig. 5 | REC curves for extremophilic variant screening.**

Regression Error Characteristic (REC) curves quantifying Screening Success Rate against absolute error tolerance for **a**, hyperstable variants ($T_m$≥60°C), **b**, thermal extreme variants ($T_{opt}$≤30°C or ≥50°C), and **c**, extreme pH variants ($pH_{opt}$≤6 or ≥8). The solid red line represents the EnzOracle model, alongside various baseline models (dashed/dotted lines). Vertical dashed grey lines indicate predefined industrial error tolerances (±5.0°C for $T_m$ and $T_{opt}$; ±0.5 for $pH_{opt}$). Red text annotations and corresponding arrows highlight EnzOracle's specific success rates at these tolerance thresholds and its absolute percentage point gain over the next best baseline.

We investigated whether the model's robustness stems from learning fundamental structure-function principles or merely overfitting to sequence homology. We evaluated predictive fidelity across varying degrees of sequence identity. At sequence identities below 30%, where sequence-based memorization typically fails, EnzOracle consistently maintained the lowest error rates. For instance, it achieved a $T_m$ RMSE of 5.143°C (vs. 5.352°C for DeepSTABp). For $T_{opt}$, EnzOracle achieves the lowest error in the remote homology range (RMSE ~11.7°C), whereas baselines like Seq2Topt and DeepET exhibit markedly higher errors (>12.5°C). The $pH_{opt}$ RMSE of 0.873 (vs. 0.895 for EpHod), confirming that the model captures universal stability constraints rather than local sequence motifs.

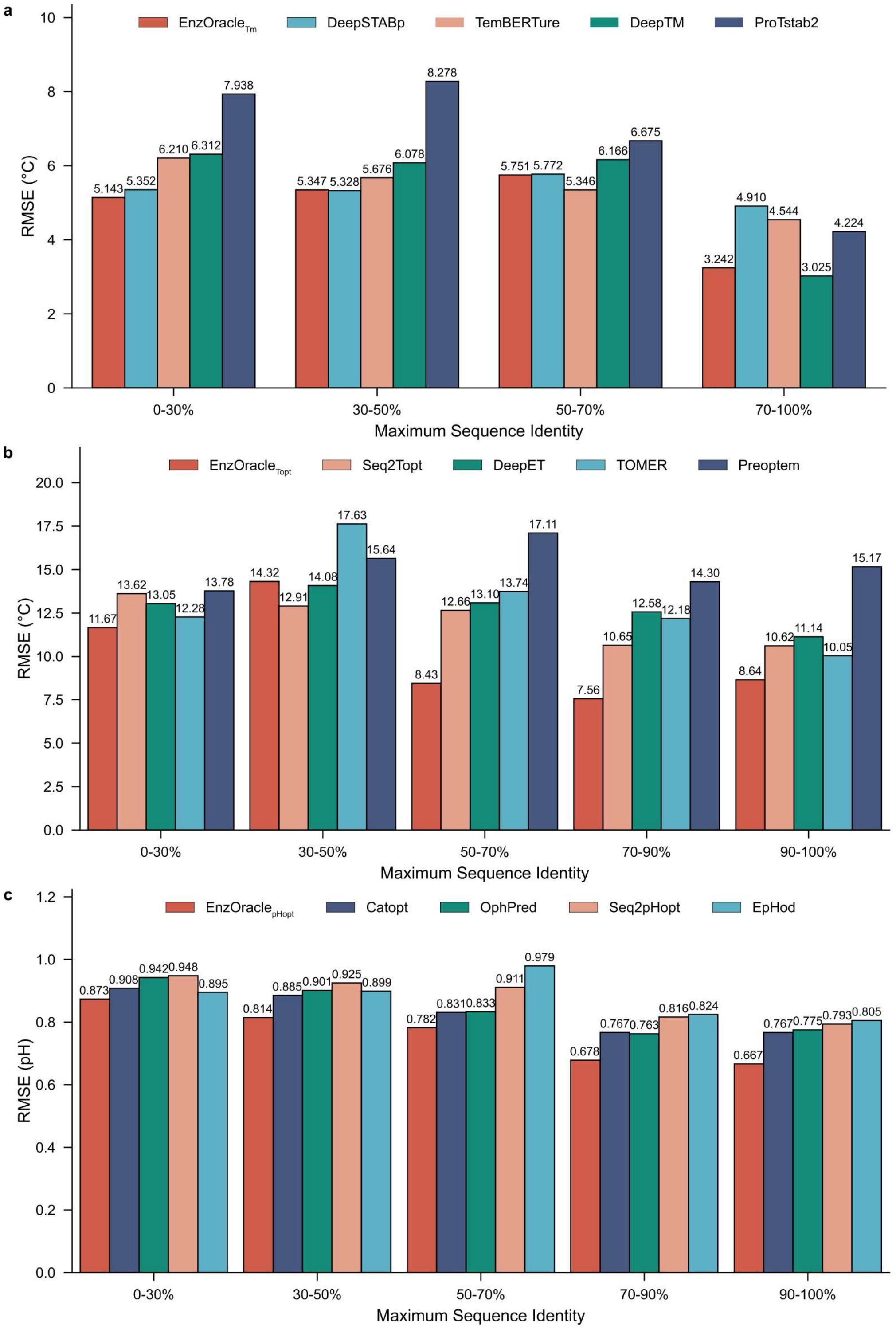


**Supplementary Fig. 6 | Generalization capability across sequence identity thresholds.**
Performance stratification by maximum sequence identity to the training set. Comparisons of RMSE for **a**, $T_m$, **b**, $T_{opt}$, and **c**, $pH_{opt}$ across binned identity ranges.

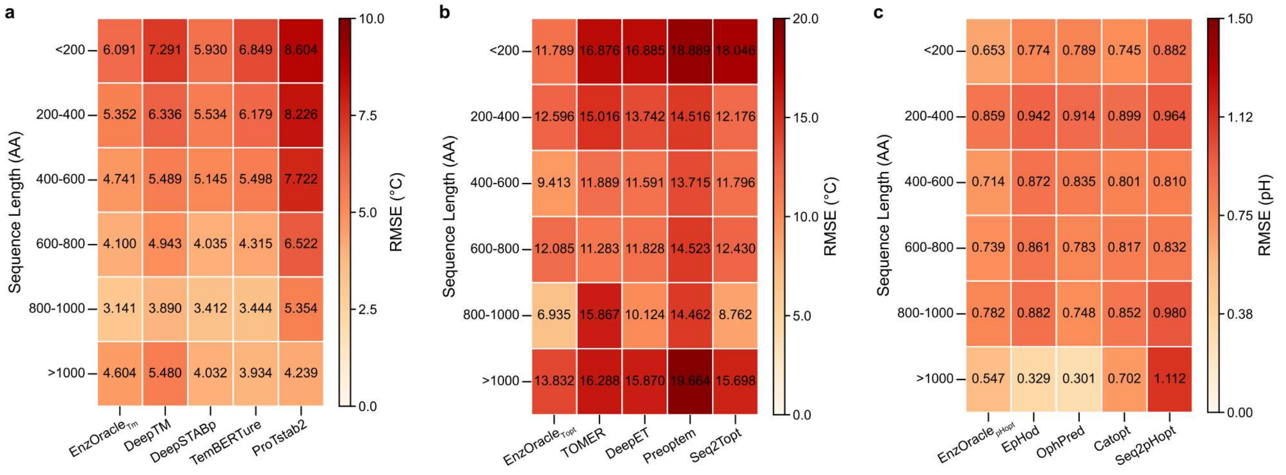


**Supplementary Fig. 7 | Robustness against sequence length variations.**

Heatmaps displaying the RMSE distribution across varying sequence lengths (<200 to >1000 AA) for **a**, $T_m$, **b**, $T_{opt}$, and **c**, $pH_{opt}$. Darker colors indicate higher prediction errors.

Analysis across Enzyme Commission (EC) categories provided the strongest evidence for functional generalization (Fig. S8). Remarkably, $pH_{opt}$ predictions maintained universal high fidelity, exhibiting the lowest error magnitudes across all seven catalytic mechanisms regardless of reaction type. However, the most substantial performance gap emerged in the challenging $T_{opt}$ task for the Ligase class (EC 6), a data-scarce category characterized by complex ATP-dependent mechanisms. Here, pure sequence-based models like Seq2Topt faltered significantly (RMSE 11.535°C). In contrast, EnzOracle achieved an RMSE of 6.054°C. This represents a 22.7% reduction in error even compared to the closest competitor DeepET (7.828°C), and a nearly 47.5% improvement over Seq2Topt.

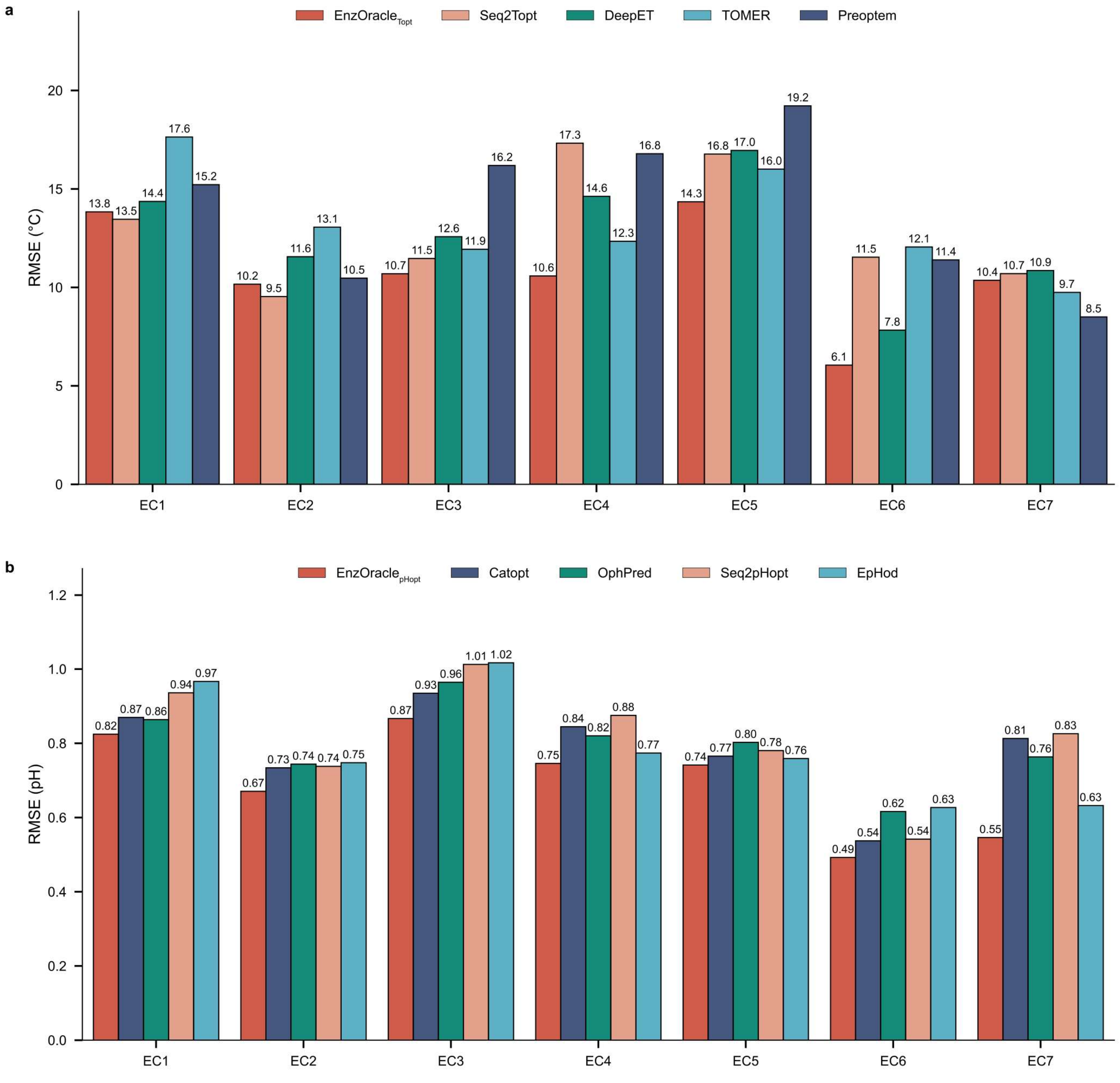


**Supplementary Fig. 8 | Broad applicability across Enzyme Commission (EC) categories.**
Benchmarking of RMSE across the seven primary EC classes for **a**, $T_{opt}$, **b**, $pH_{opt}$. Bar colors distinguish the EnzOracle model (red) from various baseline models.

Similar to the $T_{opt}$ manifold (Fig. 4a), the model organizes enzymes into topologically distinct regimes driven by data density. The landscape exhibits a clear separation between the Generalist regime (blue points) and the Specialist regime (pink points). Overlaying the expert selection markers confirms that the model establishes a discrete decision boundary: high-leverage outliers are systematically routed to the Specialist Expert (β), while the mesostable continuum is preserved for the Generalist Expert (α). This confirms that the gating network learns consistent routing strategies, effectively isolating mesostable/neutrophilic samples (routed to the Generalist) from hyperstable/extreme-pH variants (routed to the Specialist).

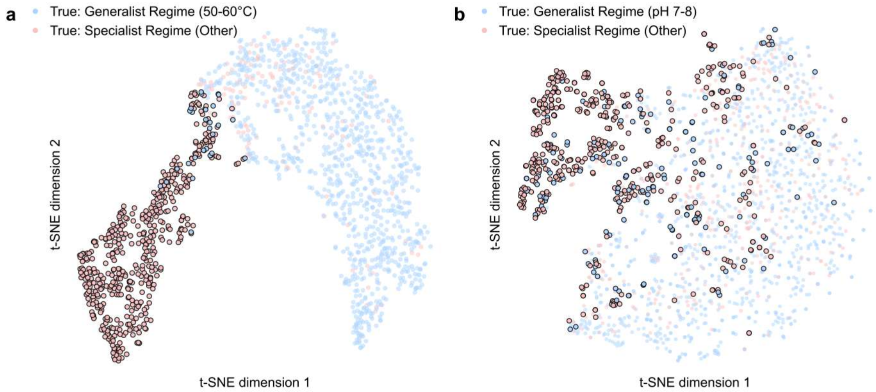


**Supplementary Fig. 9 | Latent space routing logic for stability and pH tasks.**
t-SNE visualizations of the latent feature space projected by the adaptive gating network for **a**, $T_m$ and **b**, $pH_{opt}$. Blue and pink points represent data points belonging to the Generalist Regime and Specialist Regime, respectively. Data points with a black border indicate samples assigned to the Specialist Expert by the gating network.

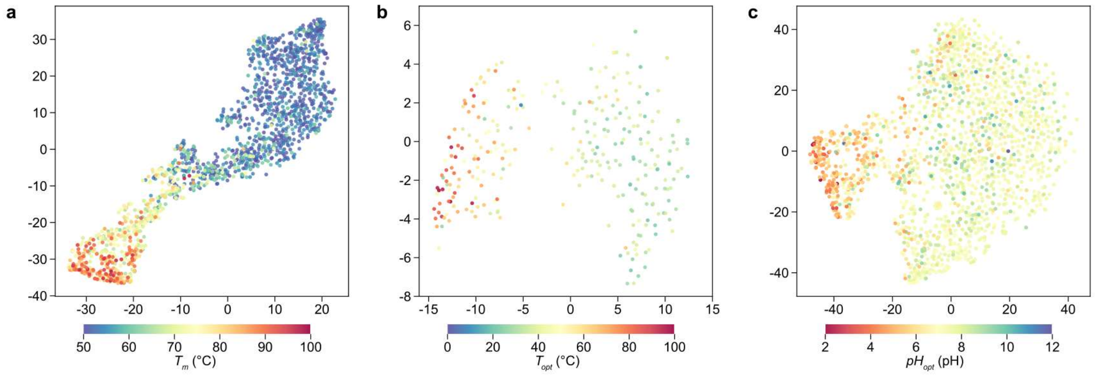


**Supplementary Fig. 10 | Emergence of continuous physicochemical manifolds.**
t-SNE visualizations of the latent feature space projected by the adaptive gating network, colored by ground-truth experimental values for **a**, $T_m$, **b**, $T_{opt}$, and **c**, $pH_{opt}$. These plots reveal the underlying continuous gradients of physical properties. The smooth transition from low (blue/cool) to high (red/hot) values confirms that the hybrid fusion of semantic (DSSE) and evolutionary (ESM-2) features results in a structured geometric space where latent spatial position directly correlates with the target physicochemical properties.

As shown in Fig. S11a, the gating dynamics for $T_m$ reveal a highly confident identification of the mesostable state, with the transition to the specialist expert occurring sharply at the 60°C threshold. This alignment with the biophysical onset of hyperstability indicates that the model has autonomously learned to treat hyperstability as a distinct physical regime requiring specialized handling. Collectively, these distinct yet structurally isomorphic gating patterns confirm that the MoE mechanism captures universal rules of extremophilic adaptation across diverse physicochemical landscapes.

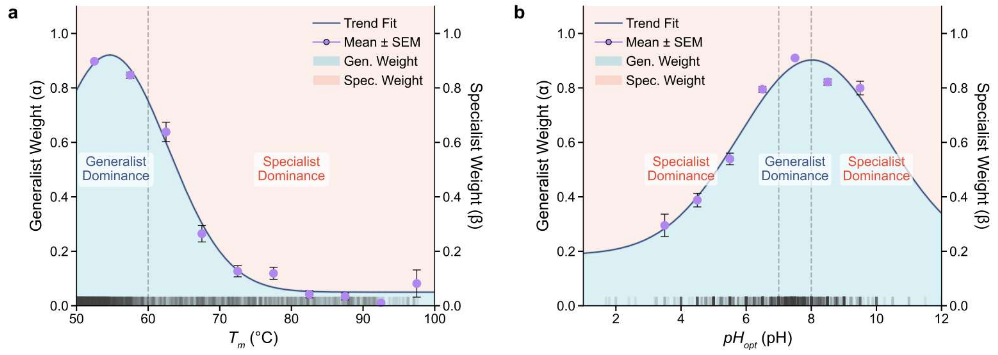


**Supplementary Fig. 11 | Adaptive gating dynamics for $T_m$ and $pH_{opt}$.**

**a**, Gating probability distribution for $T_m$. The Generalist Expert weight ($\alpha$, blue trend line) maintains a high-confidence plateau within the canonical 50-55°C range before exhibiting a sharp decay beyond 60°C, marking the transition to the Specialist Expert. **b**, Gating probability distribution for $pH_{opt}$. The gating architecture exhibits a bell-shaped curve peaking at canonical neutral conditions (pH 7-8), smoothly shifting assigning priorities to the Specialist Expert at both acidic and alkaline extremes. Visual Definitions: In both panels, purple data points represent the mean gating probability calculated within discrete intervals (5°C bins for $T_m$ and 1-unit bins for $pH_{opt}$). Error bars indicate the standard error of the mean (s.e.m.) within each bin. The background shading visually demarcates the regions of predictive dominance: light blue indicates Generalist Expert dominance ($\alpha$>0.5), while light pink indicates Specialist Expert dominance ($\alpha$<0.5). The rug plot along the x-axis visualizes the data density of the independent test set, illustrating the correlation between model confidence and actual data abundance.

For $T_m$ prediction, the gating module achieved an impressive 97.2% recall for the mesostable majority and 80.3% for the hyperstable minority (Fig. S12a). For $pH_{opt}$, the network adopted a more conservative routing strategy (Fig. S12b), achieving 86.8% recall for neutrophilic enzymes while routing only the most distinct 55.5% of acidic/alkaline variants to the specialist. This trade-off suggests that the model prioritizes precision over recall in complex pH landscapes to minimize noise injection into the specialized sub-network. This is critical not for computational savings, but for preserving the semantic purity of the Specialist Expert, preventing it from being overwhelmed by the feature dilution inherent in large-scale mesostable data rather than maximizing coverage.

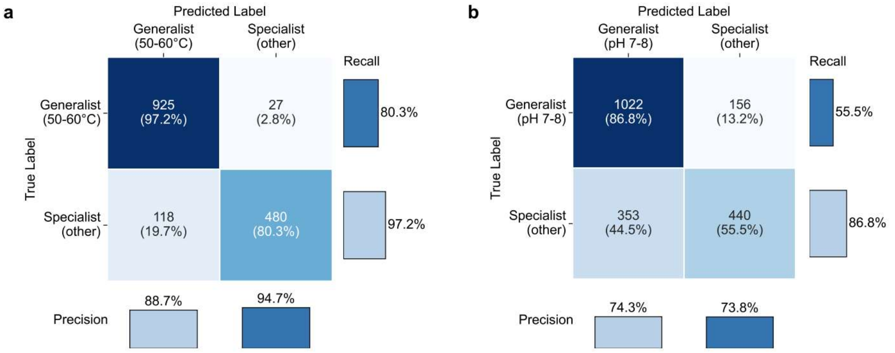


**Supplementary Fig. 12 | Discriminative reliability for $T_m$ and $pH_{opt}$.**

Extended confusion matrices evaluating the adaptive gating network's classification performance for **a**, $T_m$ and **b**, $pH_{opt}$. Central cells display absolute sequence counts and row-normalized percentages for Generalist and Specialist regime assignments, with color intensity reflecting sample density. Marginal bar charts along the right and bottom axes quantify the Recall and Precision for each class, respectively.

Decoupled RMSE trajectories demonstrating the "Rescue Effect," where the gating network dynamically selects the optimal sub-network. This complementarity is even more pronounced in $T_m$ prediction, where the dichotomy between stability mechanisms is stark (Fig. S13a). The model achieves a symmetric rescue effect: The Generalist Expert (grey dashed) prevents overfitting in the lower stability regime, achieving a 71% error reduction at 55°C compared to the Specialist Expert (red dashed). Conversely, the Specialist is critical for the hyperstable tail (80°C), reducing RMSE by 69% where the Generalist underfits. The gating mechanism leverages the Specialist Expert to reduce prediction error by 48% in acidic conditions (pH = 2) and 14% in alkaline conditions (pH = 11), effectively enveloping the minimal error bounds of both sub-networks (solid blue line). This confirms that the gating mechanism resolves the "gradient conflict" between general and extreme features. This ensures consistent fidelity across the entire physicochemical scale where monolithic models typically falter.

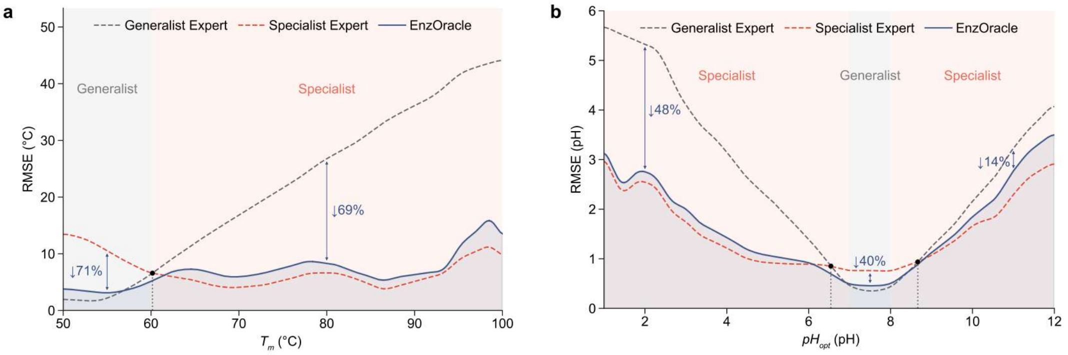


**Supplementary Fig. 13 | Complementary expert error profiles for $T_m$ and $pH_{opt}$.**

**a**, $T_m$ and **b**, $pH_{opt}$ RMSE trajectories. Curves denote the Generalist Expert (grey dashed), Specialist Expert (red dashed), and the EnzOracle model (solid blue). Background shading visually demarcates Generalist (grey) and Specialist (light pink) dominance regions. Black dots mark equilibrium crossover points, while blue arrows quantify the relative error reduction ("rescue effect") achieved by the gating mechanism.

Ablation study of architectural components (Fig. 4e). Bar charts display the $R^2$ for $T_m$ (top), $T_{opt}$ (middle), and $pH_{opt}$ (bottom). The full EnzOracle model consistently outperforms the ablated variants. Removing the pre-trained ESM-2 foundation (w/o ESM) caused the most severe performance degradation universally, effectively collapsing the model's predictive capacity; this underscores that deep evolutionary semantic information serves as the indispensable bedrock of the system. Critically, reducing the architecture to a monolithic network by removing the MoE mechanism (w/o MoE) resulted in a sharp drop in explained variance ($R^2$), particularly in tasks requiring regime separation. Using $T_{opt}$ as a representative case, the full EnzOracle model achieved an $R^2$ of 0.622, significantly outperforming the single-network baseline ($R^2$=0.541). This performance gap confirms that while deep embeddings provide the feature space, the adaptive gating mechanism is essential for resolving the variance trade-offs inherent in diverse biophysical landscapes. Furthermore, distinct from the gating and foundational layers, the exclusion of the local encoder (w/o DSSE) led to a notable performance decline (e.g., $R^2$=0.596 for $T_{opt}$). This validates the design of the Hybrid Global-Local Representation Engine (HGL-RE), demonstrating that local sequence patterns provide necessary refinement to global evolutionary priors, ensuring maximal fidelity at the amino-acid level.

For $T_m$ prediction (Fig. S14), in MetG, the classification head assigns its maximal attention (Rank 1, Score 1.0) to S427 at the C-terminal surface, identifying critical hydration sites. Conversely, the Regression Specialists systematically penetrate the protein surface to lock onto the rigid core. In PETase, the regression model identifies the "wobble" residue W185 (Rank 1, Score 1.0) and the π-stacking residue W159 (Rank 2, Score 0.985) as the primary determinants of thermal integrity. Similarly, in UV Endonuclease, the model reconstructs the internal stiffness scaffold by highlighting the Asn-ladder initiated by N107 (Rank 1, Score 1.0). This confirms that the model implicitly learns to weigh hydrophobic anchoring and hydrogen-bond networking as the physical basis of thermodynamic stability.

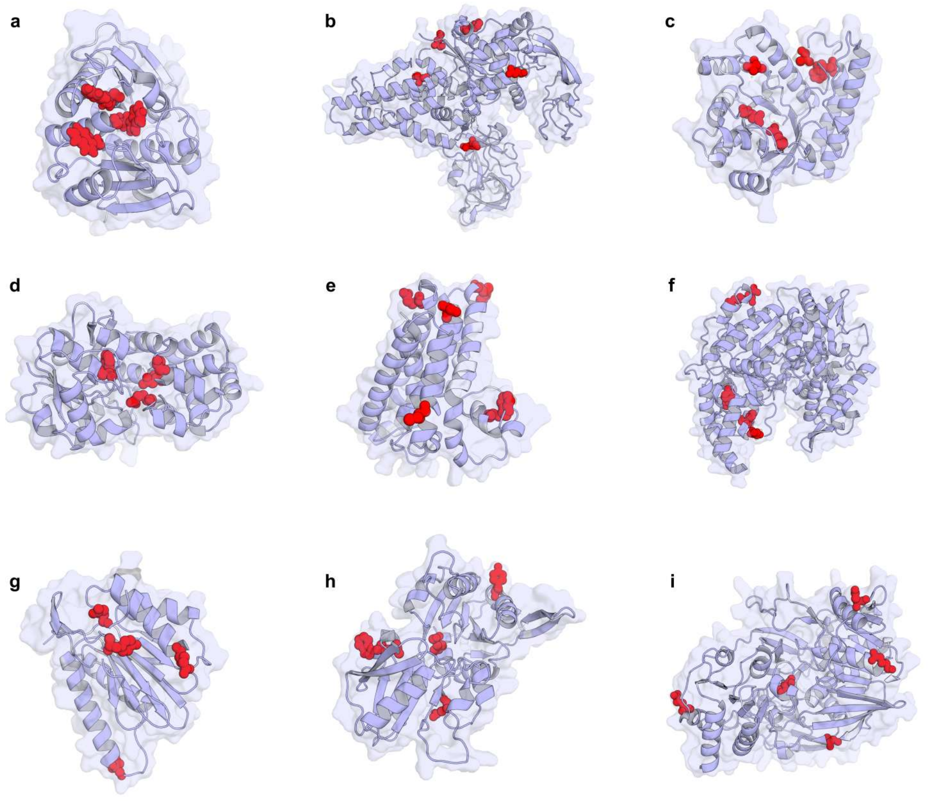


**Supplementary Fig. 14 | Visualization of structural attention weights for $T_m$ prediction.**
Light-blue regions signify residues with attention weights > 0.1, highlighting structural areas that EnzOracle emphasizes for $T_m$ predictions. Residues with weights < 0.1 are colored white. Red spheres highlight representative biologically significant residues selected from the top-20 attention candidates of the Classification, Generalist, and Specialist experts. Data in parentheses indicate UniProt accession ID (UID) and the experimental $T_m$ (in °C). **a**-**i**, *I. sakaiensis* Poly(ethylene terephthalate) hydrolase (UID: A0A0K8P6T7, $T_m$: 48.8) (**a**), Methionine-tRNA ligase (UID: Q6L1M5, $T_m$: 73.7) (**b**), Probable UV endonuclease (UID: Q746K1, $T_m$: 85.2) (**c**), Glycerol-3-phosphate dehydrogenase [NAD(P)+] (UID: P61747, $T_m$: 90.5) (**d**), Adenosylcobinamide-GDP ribazoletransferase (UID: P36561, $T_m$: 61.0) (**e**), Dihydroxyacetone phosphate acyltransferase (UID: Q545P6, $T_m$: 51.1) (**f**), Proteasome subunit $\beta$ type-4 (UID: Q7DLR9, $T_m$: 59.6) (**g**), Ribose-phosphate pyrophosphokinase 4 (UID: Q680A5, $T_m$: 58.5) (**h**) and Electron transfer flavoprotein α and $\beta$-subunit (UID: Q6L1Q7, $T_m$: 68.8) (**i**).

Distinct from the distributed attention seen in $T_m$ tasks, the $T_{opt}$ model concentrates on maintaining catalytic geometry under thermal agitation. A recurrent strategy identified by the Classification Expert is the recruitment of Tyrosine (Y) to construct aromatic cages that lock substrate-binding pockets. For instance, in Phytase (Fig. S15f), the model assigns maximal attention (Rank 1, Score 1.0) to Y377, which forms a rigid hydrophobic wall essential for stabilizing the phytate substrate at elevated temperatures. Similarly, in $\beta$-fructofuranosidase (Fig. S15g), Y293 (Rank 1, Score 1.0) is highlighted as the structural anchor of the sugar-binding cleft. Beyond hydrophobic confinement, the regression experts explicitly target mechanisms for cofactor anchoring and metallic center stabilization. In Glycerol-3-phosphate dehydrogenase (Fig. S15e), the Generalist Expert identifies Q332 (Rank 2, Score 0.934) as a critical hydrogen-bond lock for the NAD+ cofactor, preventing thermal dissociation. Furthermore, in metalloenzymes such as Deblocking aminopeptidase (Fig. S15c), the model prioritizes the zinc-coordinating residue H311 (Rank 5) alongside the C-terminal hydrophobic plug M321 (Rank 1, Score 1.0), which shields the catalytic center from solvent entropy. These patterns confirm that EnzOracle distinguishes kinetic stability, the ability to sustain catalysis, from mere structural persistence, by weighting residues that preserve the precise atomic geometry of the active site.

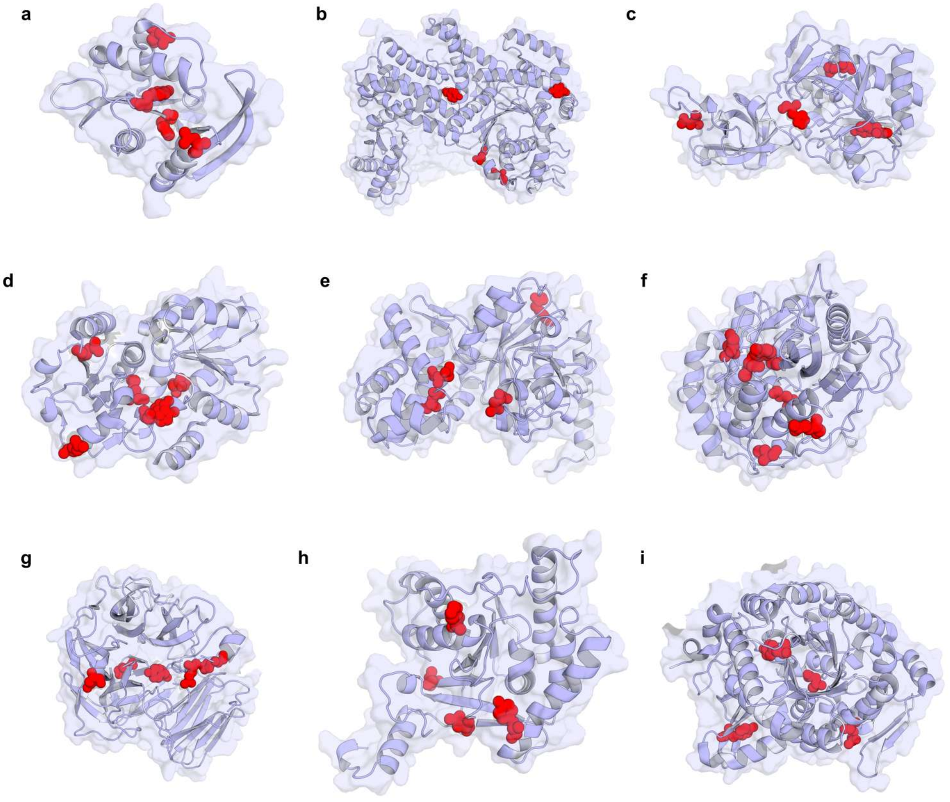


**Supplementary Figure 15 | Visualization of structural attention weights for $T_{opt}$ prediction.**

Light-blue regions signify residues with attention weights > 0.1, highlighting structural areas that EnzOracle emphasizes for $T_{opt}$ predictions. Residues with weights < 0.1 are colored white. Red spheres highlight representative biologically significant residues selected from the top-20 attention candidates of the Classification, Generalist, and Specialist experts. Data in parentheses indicate UniProt accession ID (UID) and the experimental $T_{opt}$ (in °C). **a**-**i**, *E. coli* L-methionine sulfoximine acetyltransferase (UID: Q9HUU7, $T_{opt}$: 37.0) (**a**), *T. thermophilus* Arginine-tRNA ligase (UID: O59147, $T_{opt}$: 65.0) (**b**), *P. horikoshii* Deblocking aminopeptidase (UID: Q5JHA5, $T_{opt}$: 80.0) (**c**), *A. fulgidus* Shikimate dehydrogenase (UID: P15770, $T_{opt}$: 27.0) (**d**), *C. aurantiacus* Glycerol-3-phosphate dehydrogenase (UID: A0A140JW76, $T_{opt}$: 30.0) (**e**), *E. coli* Phytase (UID: B4X9S4, $T_{opt}$: 55.5) (**f**), *T. maritima* $\beta$-fructofuranosidase (UID: Q8NMD5, $T_{opt}$: 45.0) (**g**), *P. furiosus* Diacetylchitobiose deacetylase (UID: Q6F4N1, $T_{opt}$: 72.5) (**h**) and *P. furiosus* $\beta$-galactosidase (UID: Q51723, $T_{opt}$: 95.0) (**i**).

For $pH_{opt}$ (Fig. S16), rather than simply identifying ionizable groups, the regression experts prioritize hydrophobic residues positioned to modulate the pKa of adjacent catalytic centers via hydrophobic shielding. A canonical example is observed in $\alpha$-L-arabinofuranosidase (Fig. S16a), where the model assigns near-maximal weight (Rank 3, Score 0.998) to W176. Structural analysis reveals that W176 provides a low-dielectric environment for the catalytic nucleophile E223 (Rank 17), thereby elevating its pKa for optimal activity at acidic pH. Similarly, in $\beta$-glucosidase 22 (Fig. S16e), the model identifies W206 (Rank 5, Score 0.866) shielding the catalytic proton donor E204. Beyond static pKa modulation, the model captures dynamic gating mechanisms essential for proton exchange. In GAPDH (Fig. S16b), the Generalist Expert targets the flexible S-loop hinge G173 (Rank 1, Score 1.0) and the pH-sensing residue H179 (Rank 9, Score 0.884), which acts as the catalytic base. Additionally, in Pectate lyase A (Fig. S16i), the model highlights the acidic cluster E108/E104 alongside W103 (Rank 1, Score 1.0), reflecting the pH-dependent calcium coordination required for substrate stacking. Collectively, these patterns demonstrate that EnzOracle implicitly learns to navigate the complex biophysical landscape, from surface hydration and core rigidity to electrostatic tuning, without explicit 3D supervision.

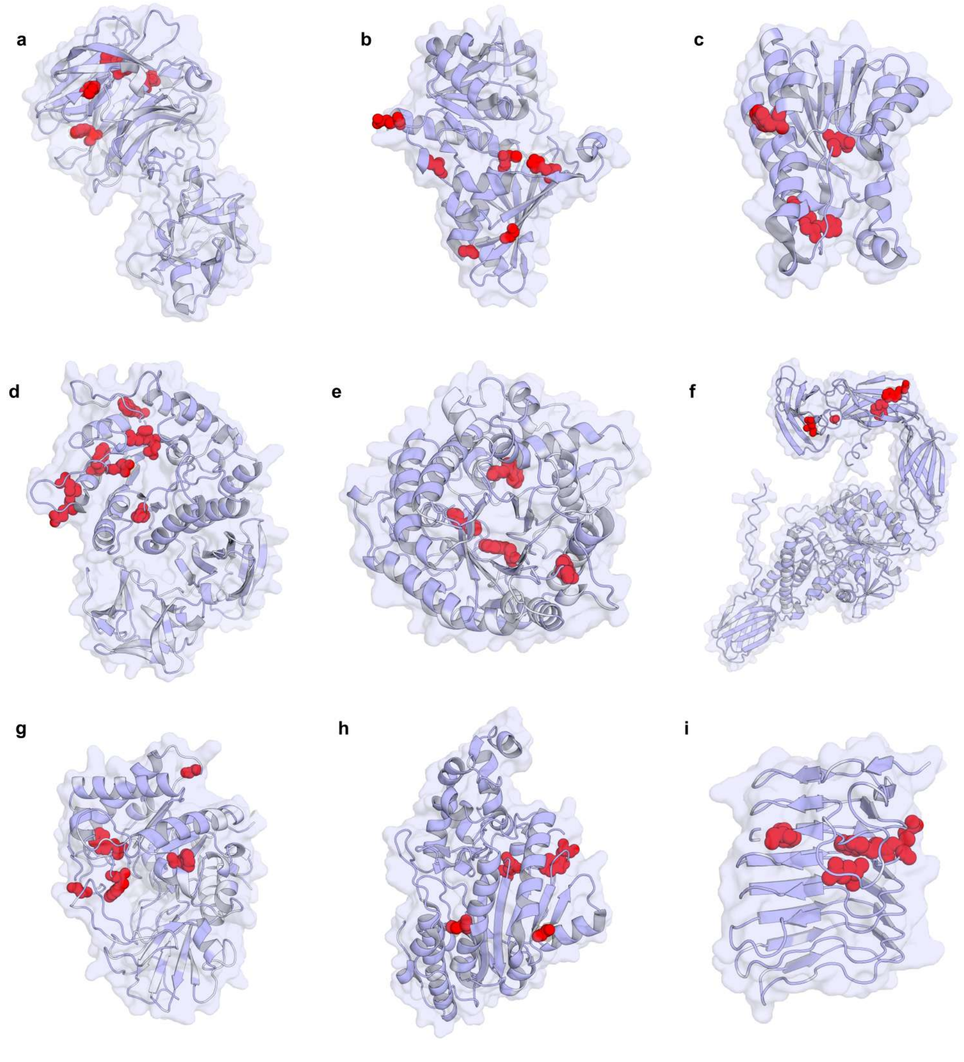

**Supplementary Figure 16 | Visualization of structural attention weights for $pH_{opt}$ prediction.**

Light-blue regions signify residues with attention weights > 0.1, highlighting structural areas that EnzOracle emphasizes for $pH_{opt}$ predictions. Residues with weights < 0.1 are colored white. Red spheres highlight representative biologically significant residues selected from the top-20 attention candidates of the Classification, Generalist, and Specialist experts. Data in parentheses indicate UniProt accession ID (UID) and the experimental $pH_{opt}$. **a**-**i**, *S. degradans* Extracellular exo-$\alpha$-(1->5)-L-arabinofuranosidase (UID: Q82P90, $pH_{opt}$: 6.0) (**a**), *B. subtilis* Glyceraldehyde-3-phosphate dehydrogenase 2 (UID: O34425, $pH_{opt}$: 8.0) (**b**), *H. sapiens* L-xylulose reductase (UID: Q7Z4W1, $pH_{opt}$: 7.0) (**c**), *G. stearothermophilus* $\alpha$-galactosidase (UID: G5D7B5, $pH_{opt}$: 2.6) (**d**), *A. thaliana* $\beta$-glucosidase 22 (UID: Q9C8Y9, $pH_{opt}$: 5.5) (**e**), *Z. mobilis* Membrane bound $\alpha$-amylase (UID: Q4J9M2, $pH_{opt}$: 3.25) (**f**), *S. marcescens* Chitinase (UID: A0A286JZ72, $pH_{opt}$: 4.5) (**g**), *H. sapiens* Creatine kinase B-type (UID: P12277, $pH_{opt}$: 9.0) (**h**) and *B. subtilis* Pectate lyase A (UID: Q9X6Z2, $pH_{opt}$: 10.0) (**i**).

Unlike the surface-focused $T_m$ model (Fig. 5a), the $T_{opt}$ attention landscape exhibits a right-skewed, bell-shaped curve. The attention weights peak significantly within the semi-exposed transition zone (30-60th percentile) before declining towards the highly disordered outer surface. Unlike $T_m$, which dictates the resistance to global structural collapse, $T_{opt}$ requires maintaining a delicate balance of conformational flexibility to enable catalytic turnover. The 30-60th RSA percentile corresponds precisely to active site pockets, substrate tunnels, and hinge loops. By focusing on these structured yet flexible regions, the $T_{opt}$ experts capture the dynamic determinants of catalysis, demonstrating that the model distinguishes between regions required for rigidity and those required for motion.

Environmental pH fluctuations primarily affect the protonation states of ionizable side chains, which interact directly with the solvent. Accordingly, the model allocates minimal attention to the hydrophobic, uncharged core (0-10th percentile), as it is insensitive to solvent proton concentration. Instead, it exhibits a broad, high-attention plateau across solvent-accessible regions, characterized by two distinct functional peaks. The first plateau (30-50th percentile) aligns with the semi-exposed active site clefts, where acid-base catalysis dictates the functional pH optimum. The global maximum, however, emerges at the highly exposed surface (70-80th percentile). This bimodal distribution accurately mirrors the dual mechanism of pH adaptation: tuning the pKa of specific catalytic residues in the pocket, while simultaneously reorganizing the global surface charge density to prevent unfolding under extreme proton concentrations.

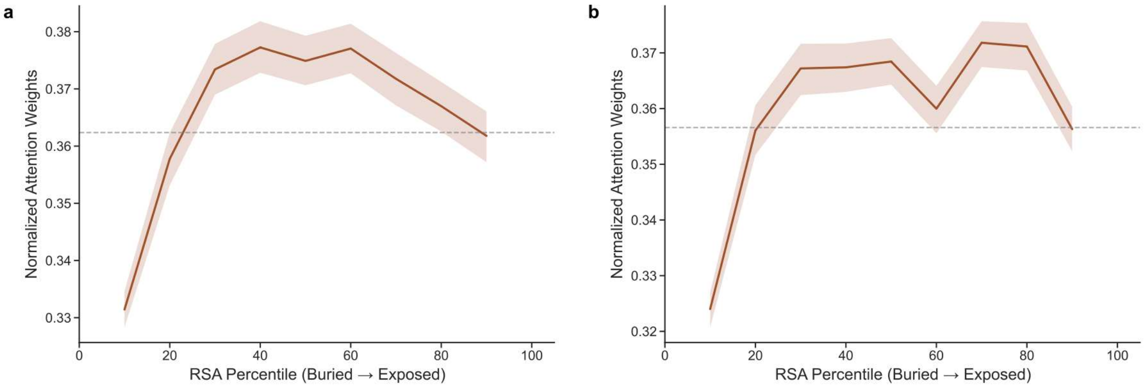


**Supplementary Fig. 17 | Structural attention landscapes for $T_{opt}$ and $pH_{opt}$.**

Gating-weighted normalized attention scores mapped across Relative Solvent Accessibility (RSA) percentiles (0 = deeply buried; 100 = highly exposed) for **a**, $T_{opt}$ and **b**, $pH_{opt}$ predictions. Solid brown lines denote the mean attention weights calculated across n=200 randomly sampled, high-confidence (pLDDT > 70) enzyme structures. This subset maintains a balanced 1:1 ratio of Generalist- and Specialist-dominated samples. Shaded regions represent 95% confidence intervals, and horizontal dashed grey lines indicate the global means.

In contrast to $T_m$ prediction, the $T_{opt}$ experts shift attention away from bulky non-polar residues and prioritize polar and catalytic residues. This demonstrates that the model actively seeks out the dynamic hydrogen-bonding networks and active-site flexibilities essential for catalytic turnover.

For $pH_{opt}$ prediction, the top-ranked residues are dominated by physiological pH sensors with titratable side chains and pH-independent aromatic structural anchors. This indicates the model captures the fundamental biophysical determinants of acid-base catalysis and electrostatic shielding.

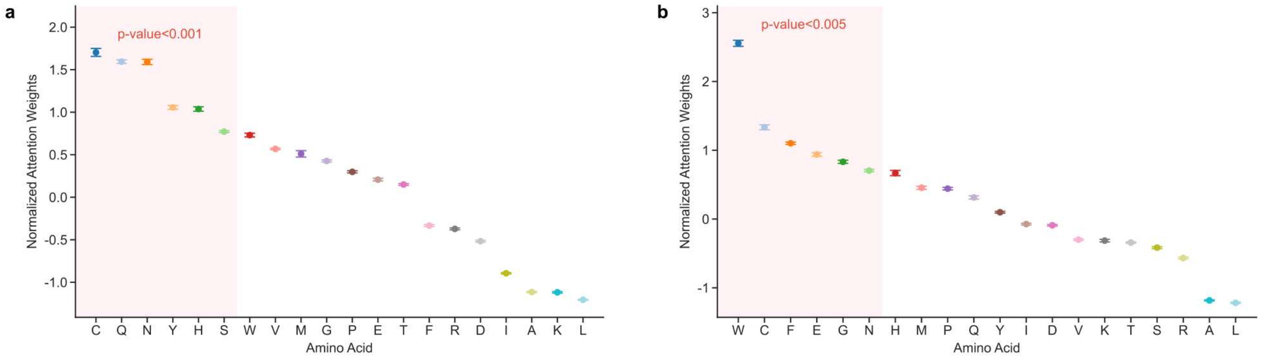


**Supplementary Fig. 18 | Global amino acid attention rankings for $T_{opt}$ and $pH_{opt}$.**

Z-score normalized, gating-weighted attention weights across the 20 standard amino acids for **a**, $T_{opt}$ and **b**, $pH_{opt}$ predictions. Data points denote mean attention weights calculated from the same subset of n = 200 randomly sampled test set enzymes used in figure. 5a. Error bars indicate the standard error of the mean (s.e.m.). Pink shaded regions highlight the top-ranked residues receiving the highest attention. Red text denotes the statistical significance for the enrichment of these highly attended subsets (two-sided Welch's t-test).

Dynamic attention shifts for $T_{opt}$ adaptation. Across psychrophilic (blue), moderate thermophilic (green), and thermophilic (red) enzymes, thermophilic adaptation is characterized by a massive decrease in attention toward rigidifying Basic residues and a highly significant peak in Polar residues. This captures the stability-activity trade-off, preventing the active site from being overly rigidified at elevated temperatures.

Dynamic attention shifts for $pH_{opt}$ adaptation. For acidophilic adaptation (high $H^+$ environments), the model massively upweights Acidic residues while severely penalizing Basic residues to prevent deleterious electrostatic repulsion. This logic is inverted for alkaliphiles, confirming that the model dynamically recalibrates its electrostatic focus according to environmental proton availability.

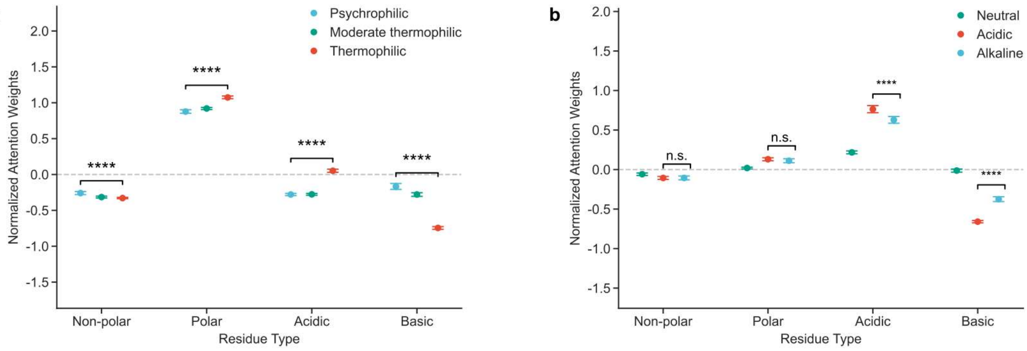


**Supplementary Fig. 19 | Environment-conditioned attention allocation grouped by residue physicochemical properties.**

Normalized attention weights across four residue types for **a**, $T_{opt}$ and **b**, $pH_{opt}$ predictions. For $T_{opt}$, enzymes are grouped into psychrophilic (< 30°C, blue), moderate thermophilic (30 − 50°C, green), and thermophilic (> 50°C, red) categories. For $pH_{opt}$, enzymes are grouped into acidic (pH < 6, red), neutral (pH 6-8, green), and alkaline (pH > 8, light blue) categories. Data points denote mean attention weights with error bars indicating the standard error of the mean (s.e.m.). Asterisks denote statistical significance across the groups (****P < 0.0001; n.s., not significant; two-sided Welch's t-test).

It is important to note that while biological functional hotspots do not necessarily occupy the absolute global attention maxima (reflecting the multi-factorial structural complexity required to maintain global protein stability) they consistently align with prominent local attention spikes. For instance, in the $T_m$ prediction of the representative enzyme (UniProt ID: P61747), the model accurately highlights the primary catalytic proton acceptor (residue 184) alongside essential cofactor and substrate binding pocket residues (e.g., residues 11, 101, 130, and 247, which coordinate NADPH and sn-glycerol 3-phosphate)(Fig. 5d). This precise structural targeting is highly consistent across different prediction tasks. When evaluating the $T_{opt}$ attention landscape of acetyltransferase Q9HUU7 (Fig. S20a), the model reveals significant attention elevations precisely localized at substrate-binding interfaces (residues 75 and 85) and critical acetyl-CoA anchoring sites (residues 88, 101, and 127). Furthermore, mapping the $pH_{opt}$ expert's attention onto creatine kinase P12277 (Fig. S20b) demonstrates accurate structural localization at the essential ATP- and creatine-binding pockets (e.g., capturing the localized regions around residues 130, 232, 285, and 320).

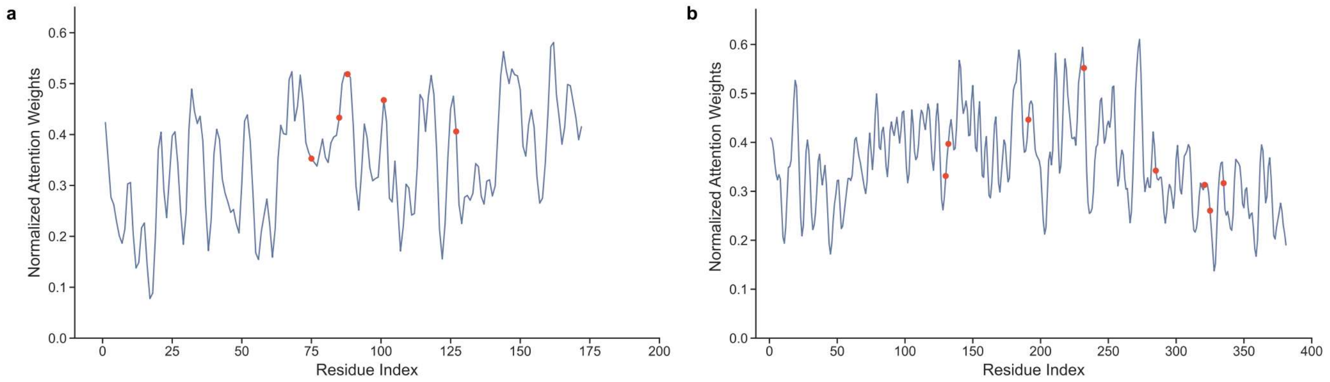


**Supplementary Fig. 20 | 1D sequence attention mapping for functional hotspots.**

Normalized attention weights of residues in the primary structures of two example enzymes. **a**, Attention profile for an acetyltransferase (UniProt ID: Q9HUU7) in $T_{opt}$ prediction. Critical substrate-binding interfaces and acetyl-CoA anchoring sites (e.g., residues 75, 85, 88, 101, and 127) are indicated by red dots. b, Attention profile for a creatine kinase (UniProt ID: P12277) in $pH_{opt}$ prediction. Essential ATP- and creatine-binding pockets (e.g., residues 130, 232, 285, and 320) are similarly indicated by red dots.

**Orthogonal molecular dynamics validation links EnzOracle-derived attention hotspots to trait-specific biophysical mechanisms**

To determine whether the structural determinants identified by EnzOracle correspond to physically meaningful molecular behavior, we performed atomistic molecular dynamics (MD) simulations across three enzyme systems representing distinct adaptive regimes. Whereas the attention-based analysis highlighted residues potentially associated with functional adaptation, MD simulations provide an orthogonal, physics-based framework to evaluate whether these sequence-derived features are linked to structural stability, residue-level fluctuations, interaction persistence, catalytic preorganization, and electrostatic organization.

We projected EnzOracle-derived attention scores onto the MD-derived enzyme structures, highlighting attention hotspots for subsequent dynamic analysis. In particular, we used MD simulations as an orthogonal, physics-based validation approach to assess whether the sequence-derived features identified by EnzOracle correspond to structurally meaningful residues. This methodology ensures that the predictive performance of the model reflects causal determinants of enzyme stability, flexibility, and electrostatic organization, which are essential for catalytic efficiency.

For the $T_{opt}$ analysis, we selected three PET-degrading enzymes: wild-type *Is*PETase, FAST-PETase, and HotPETase. These enzymes provide a suitable comparative system because they share a conserved catalytic architecture and substrate class, but differ in thermal robustness and catalytic performance. MD simulations across distinct thermal regimes revealed temperature-dependent structural responses among the three enzymes. At elevated temperatures, FAST-PETase and HotPETase rapidly converged to relatively stable conformational ensembles with backbone RMSD values of approximately 1.5Å, whereas wild-type *Is*PETase stabilized at higher RMSD values of approximately 2.1-2.4Å. These results indicate that the engineered variants better maintain their global fold under thermal perturbation, consistent with enhanced thermal resistance relative to wild-type IsPETase (Fig. S21).

PET hydrolysis proceeds through nucleophilic attack of the catalytic Ser-Oγ atom on the carbonyl carbon of the scissile ester bond, generating a tetrahedral intermediate stabilized by the oxyanion hole (Fig. 6a)[1,2]. Representative enzyme-substrate complexes extracted from 500 ns simulations reveal a conserved binding mode in which one PET repeat unit occupies the catalytic +1 subsite, flanked by upstream (-1) and downstream (+2) units (Fig. 6b). Because productive acylation requires both appropriate proximity and attack geometry, we quantified the prereaction-reactive state (PRS) using the Ser-Oγ···C distance and the corresponding Bürgi-Dunitz angle[3]. Productive near-attack conformations were defined by a nucleophile-electrophile distance d(C---Oγ)≤3.5Å and an approach angle of approximately 100-110° (Fig. 6c). PRS distance distributions revealed clear differences among the three enzymes. Wild-type *Is*PETase displayed a broad distribution extending beyond 4.0Å, indicating frequent excursions away from productive attack geometries. In contrast, FAST-PETase and HotPETase maintained narrower distributions centered around approximately 3.3Å, even at 350 K (Fig. 6d). These results suggest that the engineered variants are better able to preserve catalytically competent near-attack conformations under thermal stress. Notably, the regions assigned high attention weights by EnzOracle overlapped with residues contributing to this catalytic preorganization, supporting the interpretation that the model prioritizes structural determinants relevant to optimal-temperature function rather than merely sequence-level correlations.

Hydrogen-bond occupancy analysis of the oxyanion hole further supported this conclusion. At 330 K, FAST-PETase and HotPETase exhibited higher hydrogen-bond occupancies and shorter donor-acceptor distances than wild-type IsPETase, indicating more persistent stabilization of the developing tetrahedral intermediate (Fig. 6e, f). Because oxyanion-hole stability is essential for efficient acylation, these results provide a mechanistic link between thermal robustness, catalytic geometry, and model-derived residue importance. To further characterize substrate-enzyme interactions, we analyzed noncovalent interaction patterns in the enzyme-PET complexes using the independent gradient model based on Hirshfeld partitioning (IGMH). Both FAST-PETase and HotPETase exhibited extended interaction isosurfaces between the active-site pocket and the scissile ester bond, indicating enhanced noncovalent stabilization of the reactive substrate configuration (Fig. 6g). This observation is consistent with the PRS and hydrogen-bond analyses, suggesting that engineered PETase variants maintain a more organized catalytic environment at elevated temperature.

To complement the MD-derived conformational analysis, we optimized transition-state structures for the acylation step using a theozyme-based DFT cluster model. The optimized transition structures displayed conserved geometric features across systems, with Ser-Oγ···C distances of 1.97Å for HotPETase and 2.02Å for FAST-PETase (Fig. 6h). Although these geometric differences are subtle, they were associated with substantial differences in reaction energetics. Intrinsic reaction coordinate (IRC) analysis showed that HotPETase and FAST-PETase exhibit acylation barriers of 15.5 and 19.3 kcal/mol, respectively, indicating a more favorable acylation pathway in HotPETase.

To dissect the energetic origin of these differences, we performed distortion/interaction activation strain analysis (DIAS) using the same theozyme-based DFT cluster framework. In this analysis, the activation energy is decomposed into a distortion component, corresponding to the energetic cost required to deform the enzyme and substrate fragments toward their transition-state geometries, and an interaction component, corresponding to stabilizing interactions between the distorted fragments at the transition state. The DIAS results indicated that differences in transition-state stabilization arise primarily from more favorable interaction energies rather than from geometric distortion alone (Fig. 6i). This energetic trend is consistent with the IGMH analysis, which showed more extensive stabilizing noncovalent interactions in the engineered PETase variants. We further examined the behavior of W185, a residue implicated in PET recognition and frequently associated with active-site plasticity in PETases. In both FAST-PETase and HotPETase, W185 maintained persistent π-stacking interactions with the aromatic ring of PET, together with hydrophobic contacts involving adjacent substrate moieties. Structural superposition of MD-derived conformational ensembles showed that W185 retains sufficient local flexibility to accommodate PET while avoiding excessive disorder of the binding pocket (Fig. 6j). This balance between substrate-adaptive mobility and active-site stability provides a plausible dynamic basis for maintaining catalytic efficiency at elevated temperatures. The high attention assigned by EnzOracle to this region is therefore consistent with its role as a local dynamic hub linking substrate recognition, catalytic preorganization, and thermal adaptation.

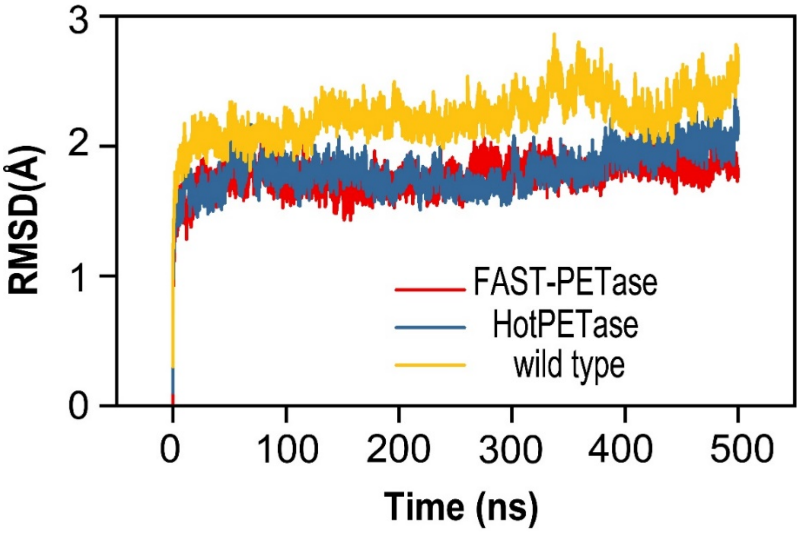


**Supplementary Fig. 21 | High-temperature RMSD analysis (350 K).**

Cα RMSD values were monitored for all three enzyme-substrate complexes under 350 K simulation conditions. The trajectories demonstrate that HotPETase and FAST-PETase, maintains significantly lower backbone deviation relative to wild type, consistent with its superior thermal stability and preservation of catalytic geometry at elevated temperature.

To validate determinants associated with melting temperature, we selected Thermus thermophilus UV damage endonuclease (UVDE) as a representative thermostable enzyme system. In contrast to $T_{opt}$, which depends strongly on local catalytic flexibility and the maintenance of productive conformations, $T_m$ primarily reflects resistance to global thermal unfolding. We therefore used UVDE to test whether residues prioritized by EnzOracle for $T_m$ prediction correspond to structurally constrained elements that stabilize the protein scaffold.

Model attribution identified N107 as a high-importance residue located within a densely packed internal region of the UVDE scaffold. To evaluate its structural role, we constructed an N107A variant in silico and compared its dynamics with the wild-type enzyme using atomistic MD simulations based on the crystal structure of UVDE. This paired wild-type/mutant design allowed us to directly assess whether the model-prioritized residue contributes to local rigidity and interaction-network persistence.

Structural analysis revealed that N107 participates in a persistent hydrogen-bonding network linking neighboring secondary-structure elements within the protein interior. The ND2 atom of N107 forms stabilizing interactions with G145, whereas the OD1 atom engages in a backbone-mediated O-H···O hydrogen bond with Y105. These interactions remained stable throughout the wild-type trajectories, with donor-acceptor distances of approximately 3.1-3.4Å and an average occupancy of approximately 63% (Fig. S22). This long-lived interaction network appears to impose a local structural constraint that helps maintain core packing geometry (Fig. S23). Disruption of this network in the N107A mutant weakened local hydrogen-bonding interactions and increased conformational flexibility in the surrounding region (Fig. S22b, c). IGMH analysis further indicated a loss of stabilizing noncovalent interaction density in the mutant relative to the wild type. These results support the interpretation that N107 acts as a rigidity-anchoring residue within the UVDE scaffold.

To evaluate whether EnzOracle captures determinants of pH adaptation, we performed MD simulations on an alkaline pectate lyase under protonation regimes corresponding to pH 7.0 and pH 10.0. Protonation states were assigned using PROPKA, enabling approximate reconstruction of pH-dependent electrostatic responses.

Model attribution analysis showed that residues assigned high importance by EnzOracle are enriched around the catalytic cleft and adjacent electrostatic interaction network, rather than being distributed primarily within the global structural scaffold. This localization suggests that $pH_{opt}$ prediction is driven largely by local charge organization and electrostatic regulation of the active site, rather than by fold-level stabilization alone. Consistent with this hypothesis, simulations under alkaline conditions revealed pronounced reorganization of interaction networks surrounding the catalytic groove (Fig.S22d). At pH 10.0, EnzOracle-prioritized residues exhibited increased salt-bridge persistence, reduced local loop fluctuations, and enhanced stabilization of the substrate-binding cleft relative to neutral conditions. The enzyme adopts a canonical parallel β-helix fold, with an extended surface cleft coordinated by $Ca^{2+}$ ions and acidic residues. Under near-neutral conditions, a primary $Ca^{2+}$ ion stabilizes the interface between the β-helix core and loop regions. Under alkaline conditions, simulations revealed recruitment of a second $Ca^{2+}$ ion within the catalytic cleft, coordinated by acidic residues including D63, E83, and D84. This additional coordination enhances the Coulombic interaction network and stabilizes the active-site environment (Fig. S22f). Radial distribution function analysis further showed that alkaline conditions induce reorganization of solvent structure surrounding the catalytic $Ca^{2+}$ ions. The first hydration shell becomes more compact (peak ~2.7Å vs. ~2.9Å at neutral pH), indicating a more ordered hydration environment. This suggests that solvent organization contributes to stabilization of metal coordination and catalytic geometry under alkaline conditions.

Together, these results indicate that pH adaptation arises primarily from redistribution of electrostatic interactions rather than changes in protonation states alone. EnzOracle-prioritized residues form persistent interaction networks that buffer electrostatic perturbations, mapping sequence-derived importance to pH-dependent structural behavior.

Thus, MD simulations serve here as a core evidential layer for evaluating the mechanistic validity of EnzOracle predictions. Rather than functioning as a supplementary visualization tool, trait-resolved MD provides a falsifiable framework for assessing whether model explanations are grounded in molecular mechanism. The observed agreement between model-derived attention hotspots and simulation-derived structural responses supports the view that EnzOracle captures transferable biophysical principles governing enzyme adaptation across thermal, catalytic, and pH-dependent regimes.

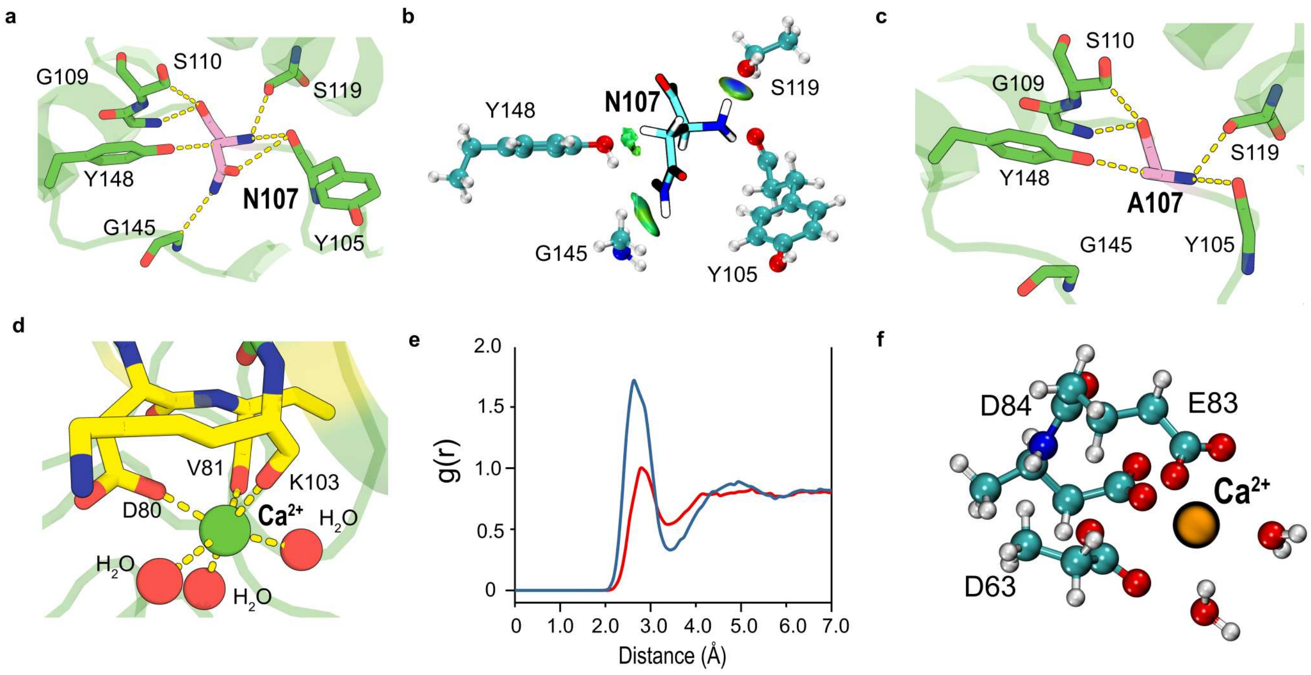


**Supplementary Fig. 22 | MD validation of melting-temperature and pH-adaptation determinants in UVDE and PelA.** **a**, Representative structure of wild-type Thermus thermophilus UVDE during MD simulation, highlighting the N107-centered interaction network identified by EnzOracle attribution. **b**, Hydrogen-bond and noncovalent interaction analysis of the N107 local environment in wild-type UVDE. IGMH isosurfaces indicate persistent stabilizing interactions within the protein core. **c**. Representative structure of the N107A mutant during MD simulation, showing disruption of the local interaction network. **d**, pH-dependent interaction-network analysis of PelA, showing enhanced salt-bridge persistence and reduced catalytic-cleft fluctuations under alkaline conditions. **e**, Radial distribution function analysis of water molecules surrounding catalytic $Ca^{2+}$ ions under neutral and alkaline conditions, showing formation of a more compact hydration shell at pH 10.0. **f**, Binding mode of the second $Ca^{2+}$ ion recruited within the PelA catalytic cleft at pH 10.0, coordinated by acidic residues including D63, E83, and D84.

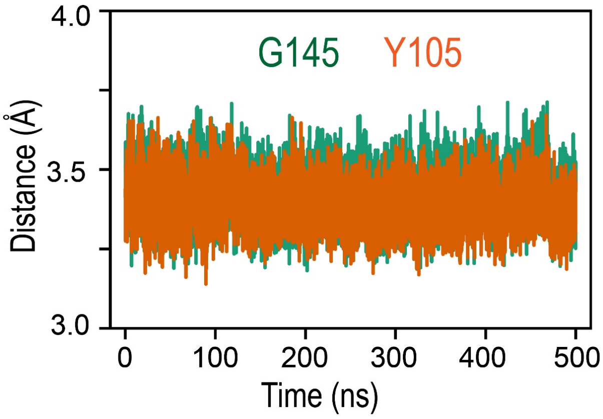


**Supplementary Fig. 23 | Analysis of the distance change between residues and Ca2+ centers during MD simulation.**

**Supplementary Table 1 | Task-specific architectural hyperparameters for the decoupled expert networks.** (Note: All nine sub-networks share the identical pre-trained ESM-2 foundation (esm2_t33_650M_UR50D) and the core Deep Semantic Sequence Encoder ( $d_{model}$=128, h=4, $L$=3 ). Only the downstream fusion and extraction parameters listed below were specifically optimized for each expert.)

| Prediction Task | Sub-network Module | Extra Global Encoder Layers | Projection Dimension | CNN Output Channels |
|---|---|---|---|---|
| | Gating Network | 3 | 512 | 512 |
| $T_m$ | Generalist Expert | 1 | 128 | 64 |
| | Specialist Expert | 3 | 512 | 512 |
| | Gating Network | 1 | 128 | 128 |
| $T_{opt}$ | Generalist Expert | 3 | 512 | 512 |
| | Specialist Expert | 3 | 512 | 512 |
| | Gating Network | 3 | 512 | 512 |
| $pH_{opt}$ | Generalist Expert | 1 | 256 | 512 |
| | Specialist Expert | 3 | 512 | 512 |

**Supplementary Table 2 | Task-specific training hyperparameters and optimization objectives for the decoupled expert networks.** All sub-networks were trained independently on NVIDIA RTX 3090 GPUs utilizing the AdamW optimizer. Hyperparameters were empirically optimized for each specific biophysical regime.

| Prediction Task | Sub-network Module | Loss Function | Learning Rate (LR) | Weight Decay | Effective Batch Size | Max Epochs | Early Stopping Patience | LR Scheduler (Warmup) |
|---|---|---|---|---|---|---|---|---|
| $T_m$ | Gating Network | BCELoss | 2e-5 | 1e-5 | 16 | 500 | 18 | ReduceLROnPlateau |
| | Generalist Expert | Weighted SmoothL1 (0.1) | 1e-4 | 1e-3 | 32 | 200 | 20 | CosineAnnealingLR |
| | Specialist Expert | RegFocalLoss | 2e-5 | 1e-5 | 16 | 500 | 18 | ReduceLROnPlateau |
| $T_{opt}$ | Gating Network | LabelSmoothing BCE (0.1) | 5e-5 | 1e-2 | 64 (16×4)* | 500 | 12 | CosineAnnealing WarmRestarts |
| | Generalist Expert | RegFocalLoss | 2e-5 | 1e-5 | 16 | 500 | 18 | ReduceLROnPlateau |
| | Specialist Expert | RegFocalLoss | 2e-5 | 1e-5 | 16 | 500 | 18 | ReduceLROnPlateau |
| $pH_{opt}$ | Gating Network | BCEWithLogitsLoss | 1e-4 | 1e-2 | 32 (8×4)* | 500 | 18 | Cosine Annealing (2% Warmup) |
| | Generalist Expert | MSELoss | 2e-5 | 1e-2 | 32 (8×4)* | 200 | 18 | Cosine Annealing (5% Warmup) |
| | Specialist Expert | WeightedMSELoss | 1e-4 | 1e-5 | 32 (8×4)* | 300 | 30 | Cosine Annealing (5% Warmup) |

* Note: Effective batch size calculated as (Physical Mini-batch Size × Gradient Accumulation Steps).

**Supplementary Note: Formulation of Objective Functions**

**1. Binary Cross-Entropy (BCE) Loss**

In the $T_m$ prediction task, we employed the standard BCE loss. The BCE loss quantifies the divergence between the predicted probability distribution and the discrete ground-truth environmental labels. For a given mini-batch of size $N$, the objective function is mathematically formulated as:

$$\mathscr{L}_{BCE} = -\frac{1}{N}\sum_{i=1}^{N} [y_i \log(\hat{y}_i) + (1-y_i)\log(1-\hat{y}_i)] \quad (1)$$

where $y_i \in \{0,1\}$ denotes the ground-truth binary label for the $i$-th sequence (assigned as 1 for the data-dense canonical regime and 0 for the extremophilic tail), and $\hat{y}_i \in (0,1)$ represents the continuous probability scalar output by the discriminator's sigmoid activation.

**2. Binary Cross-Entropy with Logits Loss incorporating Label Smoothing**

In the $T_{opt}$ prediction task, we implemented a Binary Cross-Entropy with Logits Loss augmented with label smoothing. Standard binary classification relies on hard environmental labels ($y_i \in \{0,1\}$), which can compel the network to learn excessively large logit values, often resulting in severe model overconfidence and poor generalization near the physicochemical classification boundaries.

To mitigate this, label smoothing relaxes the hard ground-truth targets into soft probabilities. Given a smoothing factor $\epsilon$ (empirically set to 0.1 in our configuration), the smoothed target $y_{new}^{(i)}$ for the $i$-th sequence is transformed as follows:

$$y_{new}^{(i)} = y_i(1-\epsilon) + 0.5\epsilon \quad (2)$$

Consequently, positive labels ($y_i = 1$) are decayed to 0.95, while negative labels ($y_i = 0$) are smoothed to 0.05. The final objective function computes the log-loss between these soft targets and the unnormalized logits ($x_i$) while internally applying the sigmoid activation ($\sigma$) for numerical stability:

$$\mathscr{L}_{LS-BCE} = -\frac{1}{N}\sum_{i=1}^{N} \left[y_{new}^{(i)} \log(\sigma(x_i)) + (1-y_{new}^{(i)})\log(1-\sigma(x_i))\right] \quad (3)$$

By preventing the discriminator from assigning absolute absolute certainty (probabilities of exactly 0 or 1), this smoothed formulation yields a highly calibrated, continuous gating scalar. This calibration is essential for ensuring robust interpolation between the Generalist and Specialist expert networks during late fusion.

**3. Binary Cross-Entropy with Logits Loss (BCEWithLogitsLoss)**

In the $pH_{opt}$ prediction task, we utilized the Binary Cross-Entropy with Logits Loss. Unlike the standard BCE loss, this objective function directly operates on the unnormalized raw logit outputs from the discriminator network, fusing the Sigmoid activation layer and the BCE loss into a unified mathematical operation. For a mini-batch of size $N$, the loss is defined as:

$$\mathscr{L}_{BCEWithLogits} = -\frac{1}{N}\sum_{i=1}^{N} [y_i \log(\sigma(x_i)) + (1-y_i)\log(1-\sigma(x_i))] \quad (4)$$

where$x_i$represents the raw logit prediction for the $i$-th sequence, $y_i \in \{0,1\}$ denotes the discrete ground-truth environmental regime label, and $\sigma(x_i) = (1+\exp(-x_i))^{-1}$ is the sigmoid activation function. By integrating the sigmoid computation within the log-loss framework, this formulation inherently leverages the log-sum-exp trick. This integration ensures superior numerical stability by preventing gradient underflow or overflow, which is critical for maintaining robust and continuous probabilistic gating during the complex optimization landscape of extremophilic data.

**4. Weighted Smooth L1 Loss**

In the Generalist Expert regression task for $T_m$, we implemented a Weighted Smooth L1 Loss. Standard regression objectives are often overwhelmingly biased toward densely populated data regimes and highly sensitive to target outliers. To systematically mitigate this bias, the continuous target space was first discretized into fixed-width intervals (e.g., 5 °C bins) to reflect the thermal distribution. For an instance $i$ belonging to bin $k$, a sample-specific weight $\omega_i$ was calculated utilizing an inverse-frequency approach to penalize mispredictions on rare extreme variants:

$$\omega_i = 1 - \frac{N_k}{N_{total}} \tag{5}$$

where $N_k$ denotes the sample count within bin $k$, and $N_{total}$ is the overall number of training samples.

The final objective function integrates these data-driven weights with the Smooth L1 (Huber) loss, which combines the robustness of L1 loss against outliers with the smooth gradient characteristics of L2 loss near the optimal convergence point. For a mini-batch of size $B$, the loss is mathematically formulated as:

$$\mathcal{L}_{WeightedSmoothL1} = \frac{1}{B}\sum_{i=1}^{B} \omega_i \begin{cases} \frac{0.5(\hat{y}_i - y_i)^2}{\beta} & if\ |\hat{y}_i - y_i| < \beta \\ |\hat{y}_i - y_i| - 0.5\beta & otherwise \end{cases} \tag{6}$$

Where $\hat{y}_i$ and $y_i$ are the predicted and experimental physicochemical values, respectively, and the transition threshold parameter $\beta$ was empirically set to 0.1. This hybrid formulation ensures robust and stable gradient propagation while dynamically directing the model's attention toward the data-scarce extremophilic tails.

#### 5. Regression Focal Loss (RegFocalLoss)[4]

To rigorously address the challenge of learning from highly sparse and imbalanced extremophilic distributions, we implemented a Regression Focal Loss (RegFocalLoss). This objective function was specifically deployed for the Specialist Expert in the $T_m$ task, as well as both the Generalist and Specialist Experts in the $T_{opt}$ task. Standard Mean Squared Error (MSE) treats all samples equally, causing the model to be easily dominated by easily predictable majorities while underfitting the "hard" boundary or extreme samples.

RegFocalLoss mitigates this by dynamically scaling the squared error based on the current absolute prediction error of each sample within a batch. For a mini-batch of size $B$, let $e_i = |\hat{y}_i - y_i|$ denote the absolute error for the $i$-th sequence. A normalized, gradient-detached focal weight $\alpha_i$ is dynamically computed as:

$$\alpha_i = \frac{e_i^{\gamma}}{\sum_{j=1}^{B} e_j^{\gamma} + \epsilon} \tag{7}$$

where $\gamma$ is the focusing parameter (empirically set to 1.0) and $\epsilon = 10^{-8}$ ensures numerical stability. By strictly detaching $\alpha_i$ from the computational graph, it acts purely as a dynamic modulating scalar without disrupting the underlying gradient flows. Consequently, the network autonomously shifts its optimization focus toward instances with larger current errors (i.e., hard-to-predict variants).

Furthermore, to counteract the static, global data imbalance, this dynamic focal loss is multiplied by the inverse-frequency sample weight $\omega_i$. Consistent with the methodology described previously, $\omega_i$ was calculated via target discretization (employing 5 °C intervals for the $T_m$ dataset and 10 °C intervals for the $T_{opt}$ dataset). The finalized objective function is mathematically formulated as:

$$\mathcal{L}_{RegFocal} = \frac{1}{B}\sum_{i=1}^{B} \omega_i \cdot \alpha_i \cdot (\hat{y}_i - y_i)^2 \tag{8}$$

This dual-weighting paradigm—combining global static distribution penalties ($\omega_i$) with batch-level dynamic error focusing ($\alpha_i$)—ensures robust convergence and exceptional predictive fidelity across both broad canonical landscapes and ultra-rare thermal extremes.

#### 6. Mean Squared Error (MSE) Loss

For the Generalist Expert deployed in the $pH_{opt}$ prediction task, we utilized the MSE loss. Unlike the sparse extremophilic regimes that necessitate aggressive re-weighting or focal strategies, the canonical pH landscape (specifically defined here as $7.0 \le pH_{opt} \le 8.0$) is characterized by a high density of structurally diverse mesophilic data. Consequently, an unweighted L2 penalty is mathematically optimal for capturing the central tendency of this abundant and relatively balanced distribution.For a given mini-batch of size $B$, the MSE loss is formulated as:

$$\mathcal{L}_{MSE}=\frac{1}{B}\sum_{i=1}^{B}(\hat{y}_i-y_i)^2 \tag{9}$$

where $\hat{y}_i$ and $y_i$ denote the predicted and experimentally determined $pH_{opt}$ values for the $i$-th sequence, respectively. By quadratically penalizing prediction deviations without introducing artificial sample weights, this straightforward yet robust objective function ensures rapid, stable gradient descent. This allows the Generalist Expert to maintain exceptional predictive fidelity and smooth interpolation across the dominant mesophilic biocatalyst space, strictly avoiding over-parameterization where it is unwarranted.

**7. Weighted Mean Squared Error (WeightedMSELoss)**

For the Specialist Expert dedicated to the $pH_{opt}$ prediction task, the dataset predominantly consists of acidic and alkaline extremophiles. Unlike the canonical mesophilic regime, this extremophilic space suffers from severe internal data sparsity and highly skewed frequency disparities. To prevent the model from ignoring ultra-rare pH variants while simultaneously avoiding gradient explosion, a sophisticated Weighted Mean Squared Error (WeightedMSELoss) was implemented.

The continuous pH spectrum was first discretized into 1-unit intervals (bins). To calculate the sample-specific weight $\omega_i$ for a sequence falling into bin $k$, we employed an inverse-square-root frequency heuristic rather than a strict inverse frequency. This mathematically softens the penalty scaling, achieving a robust balance between learning minority extremophiles and preserving overall predictive accuracy. The pre-normalized weight $w_{raw,k}$ is defined as:

$$w_{raw,k}=\frac{1}{\sqrt{N_k}} \tag{10}$$

where $N_k$ represents the sequence count within bin $k$. To ensure training stability and optimize global regression metrics (such as $R^2$), the weights were subsequently normalized to maintain a global mean of 1.0. Crucially, an empirical clipping function was applied to explicitly bound the maximum weight (capped at a threshold of 6.0 in our configuration). This dynamic clipping maintains a stable penalty ratio (maximum-to-minimum weight ratio), effectively preventing aberrant outlier samples from disproportionately dominating the gradient updates.

Following the final mean-normalization, the sample-specific weight $\omega_i$ is integrated into the MSE formulation. For a mini-batch of size $B$, the objective function is mathematically formulated as:

$$\mathcal{L}_{WeightedMSE}=\frac{1}{B}\sum_{i=1}^{B}\omega_i(\hat{y}_i-y_i)^2 \tag{11}$$

where $\hat{y}_i$ and $y_i$ denote the predicted and experimental $pH_{opt}$ values, respectively. This customized objective ensures that the Specialist Expert diligently captures the physical rules governing extreme pH adaptation without sacrificing the statistical stability of the learning process.

**Supplementary Note: MD simulations**

Conformational sampling of the substrate was performed using the CREST tool based on the semiempirical GFN2-xTB quantum chemical method developed by Grimme and co-workers. To derive force field parameters for the substrate, electrostatic potential (ESP) calculations were carried out at the HF/6-31G(d) level, followed by a two-stage restrained electrostatic potential (RESP) charge fitting protocol[5]. The bonded and non-bonded parameters, including bond lengths, angles, dihedral angles, and van der Waals radii, were generated using the Antechamber module within the AmberTools suite[6]. Molecular docking calculations were performed using AutoDock Tools[7,8].

Through tleap, systems were solvated in an octahedral TIP3P water box, ensuring an external water layer thickness of at least 10Å, and sodium ions were added to neutralize the charge. The solvated systems underwent energy minimization to eliminate any atomic collisions. Subsequently, the systems were gradually heated from 0 K to 300 K over 100 ps under the Langevin thermostat, followed by equilibration in the NPT ensemble for 200 ps to achieve stable density. Long-range electrostatic interactions were handled using the Particle Mesh Ewald (PME) method.
